# Uncovering the hologenomic basis of an extraordinary plant invasion

**DOI:** 10.1101/2022.02.03.478494

**Authors:** Vanessa C. Bieker, Paul Battlay, Bent Petersen, Xin Sun, Jonathan Wilson, Jaelle C. Brealey, François Bretagnolle, Kristin Nurkowski, Chris Lee, Gregory L. Owens, Jacqueline Y. Lee, Fabian L. Kellner, Lotte van Boheeman, Shyam Gopalakrishnan, Myriam Gaudeul, Heinz Mueller-Schaerer, Gerhard Karrer, Bruno Chauvel, Yan Sun, Love Dalén, Péter Poczai, Loren H. Rieseberg, M. Thomas P. Gilbert, Kathryn A. Hodgins, Michael D. Martin

## Abstract

While invasive species are a key driver of the global biodiversity crisis, the drivers of invasiveness remain debated. To investigate the genomic basis of invasiveness in plants, we use the invasive weed *Ambrosia artemisiifolia*, introduced to Europe in the late 19^th^ century, resequencing 655 ragweed genomes, including 308 herbarium specimens collected up to 190 years ago. In introduced European populations, we report selection signatures in defense genes and lower prevalence of particular plant pathogens in the invasive range. Together with temporal changes in population structure associated with introgression from closely related *Ambrosia* species, escape from microbial enemies likely favoured the plant’s remarkable success as an invasive species.

**One-Sentence Summary:** The invasive success of European ragweed was facilitated by release from enemy microbes and inter-species hybridization.

## Main Text

The wide-scale introduction of exotic species to novel ranges around the world can be largely attributed to the nineteenth-century colonial activities of Europeans and to escalating global trade activities since the twentieth century (*1*). Invasive species are now one of the major drivers of ecological change (*2*). They threaten global biodiversity and ecosystems by outcompeting native species (*3, 4*). They also have a large economic impact, with terrestrial invasive species costing an estimated 134 billion USD in the United States alone (*5*). Attempts to stymie the rate of new introductions have failed on a global scale, as there has been no saturation in the accumulation of alien species worldwide, and the rate of new introductions may even be accelerating (*1*).

Many more species are introduced to novel ranges than become invasive. One of the fundamental questions in invasion biology is why some aliens become invasive while others fail even to establish a permanent population (*6*). Some hypotheses attempt to explain differential success of invasive species in relation to their traits. For example, certain characteristics of some plants (‘ideal weeds’) make them more prone to become invasive, including prolific production of long-lived seeds, the rapid growth of seedlings, no biological necessity for specialized pollinators, self-compatibility, and adaptations for long-distance dispersal (*7, 8*). Adaptive genetic changes are common and often important following initial introduction (*9–11*). Other hypotheses explaining species’ differential success as invaders invoke changes in ecological interactions in the introduced range, including escape from herbivores and pathogens (enemy-release hypothesis, ERH (*12, 13*)). In plants, it has been proposed that the escape from specialized enemies in the native range allows an exotic species to allocate resources from defence mechanisms towards growth and reproduction, increasing its competitiveness in the introduced range (evolution of increased competitive ability, EICA (*12, 14*)).

In addition to herbivorous animals, exotic introduction may also free plants from fungal (*15*), bacterial (*16*), oomycete (*17*), or viral (*18*) pathogens that live on their surface or within their tissues. Plants introduced to new geographic regions will also face new microbial interactions (*19*) that affect the plant’s fitness and in some cases could facilitate invasive success (*20–22*). So far, few studies have investigated the influence of microbial communities on the success of invasive plants, and never at the genomic level. Considering the important role of plant-microbe interactions in evolutionary ecology (*23*), a holistic characterization of the invasion process should also investigate the interplay of the host genome with its associated microbial metagenome (*24–27*).

To investigate the evolutionary genomic basis of plant invasion, we chose *Ambrosia artemisiifolia* (common ragweed), an extraordinarily successful noxious weed that is native to North America with a 200-year history of global introductions (*28, 29*). It is among the 100 most impactful exotic species in Europe (*30*), and has established invasive populations in >30 European countries (*31*). *A. artemisiifolia* causes increasingly negative economic and public health impacts (*32*), mostly owing to its prolific production of highly allergenic, windborne pollen (*33*). Its future success is linked to climate change; thus it is predicted to become a more serious problem in the coming decades (*34, 35*). As shown by previous studies, *A. artemisiifolia* is able to rapidly adapt to its new environment (*36–39*) and thus has great potential to expand and become invasive in more regions. Recent work estimating the potential impact of biological control with the ragweed leaf beetle (*Ophraella communa*) offers some hope for reducing ragweed’s impact in Europe (*32*), and an indication that understanding ragweed’s ecological interactions may be one key to success in slowing the plant’s invasion.

## Results

To uncover the hologenomics of invasion in this exceptionally successful invasive plant, we report a *de-novo* assembly and annotation of the nuclear genome of common ragweed with a further analysis of 655 temporally sampled individual genomes and metagenomes. Nearly 50% of these samples come from historical herbarium collections. We grouped samples into five populations based on geography and genetic clustering. We found that the main source of the introduced European invasive population is a native-range admixed genetic cluster that likely arose due to the anthropogenic activities of early European colonists in North America.

We found large temporal changes in population structure in Europe, but not in North America, with several genetic clusters being exclusive to modern Europe. All spatial groups in Europe show signals of introgression from closely related *Ambrosia* spp. In Europe, we found evidence of recent selection on genes associated with defense, plant growth, and flowering time, and differences in the presence and prevalence of plant pathogens between Europe and North America, consistent with ERH.

### De-novo assembly of nuclear genome

The initial Meraculous (*40, 41*) assembly of the short-read data from individual AA19_3_7 resulted in an assembly of length 1579.1 Mbp, composed of 93,647 scaffolds with an N_50_ of 89.7 Kbp. After filtering this initial assembly to remove haplotigs, the resulting filtered assembly consisted of 12,288 contigs with a total length of 1280.34 Mbp and an N_50_ of 101.591 Kbp. After HiRise scaffolding with the Chicago sequencing data, the final genome assembly’s length was 1258.37 Mbp, and it was composed of 12,228 scaffolds with a scaffold N_50_ of 270.6 Kbp. The repeat analysis resulted in an annotation of 30.76% of the genome sequence in interspersed repeats. Of the whole genome, 8.01% are long terminal repeat (LTR) elements, 2.01% are long interspersed nuclear elements (LINEs), 3.75% are DNA transposable elements, 0.20% are short interspersed nuclear elements (SINEs), and 16.79% are unclassified repeats. The gene annotation of the repeat-masked genome resulted in 34,066 predicted proteins. The benchmarking universal single-copy orthologs (BUSCO) (*42*) analysis of assembly completeness determined the state of 1,375 single-copy ortholog genes, finding 756 (54.9%) were complete and single-copy, 176 (12.8%) were complete and duplicated, 68 (5.0%) were fragmented, and 375 (27.3%) were missing.

### Spatio-temporal population structure

In the native North American (NA) range, samples cluster based on geography in both the PCA (Fig. 1) and admixture (Fig. 2) analysis, although genetic differentiation as measured by F_ST_ is low between these populations (Fig. 1). There are four main genetic clusters observed for *K*=9 in the native range: the light pink cluster that is the main component of NA West samples, the dark pink cluster that is the main component of samples from NA East, the light turquoise cluster that is the main component of samples from NA South and a blue cluster that is found in in the Admixed population that is located between the three extremes of the species range in North America (Fig. 2). The geographic clustering in the native range did not change substantially between historical (collected between 1830 and 1973) and modern (collected between 2009 and 2019) times, and F_ST_ values are low when comparing the same populations through time (mean F_ST_=0.008, Fig. 1). The highest F_ST_ values can be observed in comparisons with the NA South population; although it is overall the most divergent, it is more similar to the NA East than the NA West population. The South population contributed little to the Admixed population as evidenced by a clear separation of the South population from the other populations in the PCA, admixture proportions, and high F_ST_ values. The Admixed population is located between the East and West population on the PCA, connecting the two clusters and leading to a continuous distribution rather than a clear separation of the clusters. In the historical time period, it is closer to the West population, with the F_ST_ value being about twice as high between Admixed - East than between Admixed - West. In the modern time period, this difference disappears, with Fst between Admixed - East being nearly identical to Fst between Admixed – West. Based on D-statistics, we found evidence of introgression from the related species *Ambrosia trifida* in spatial groups from the West population. We also found evidence of introgression from *Ambrosia psilostachya* in spatial groups from the South and West populations based on D-statistics (Fig. S1-2) and TreeMix (Fig. S3). PSMC-based demographic reconstruction of a high-depth North American sample shows a population decline between 10^5^ and 10^4^ years ago (Fig. S4).

**Fig. 1.**
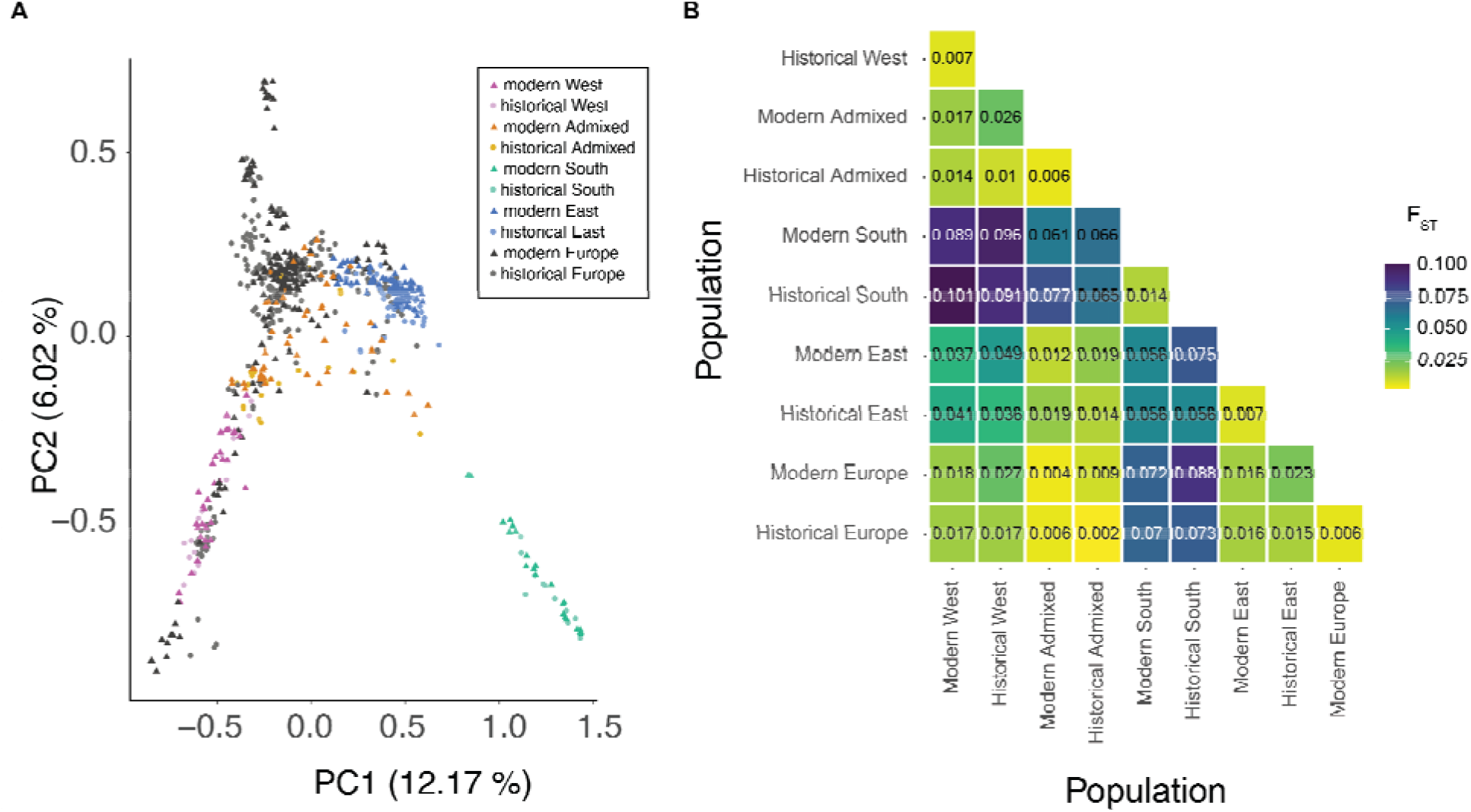
Population structure in *Ambrosia artemisiifolia*. (**A**) PCA of *A. artemisiifolia* samples. North American populations are defined based on genetic clustering and geography. Dark pink: modern West NA, light pink: historical West NA, dark orange: modern Admixed NA, light orange: historical Admixed NA, dark turquoise: modern South NA, light turquoise: historical South, dark blue: modern East NA, light blue: historical East NA, dark grey: modern Europe, light gray: historical Europe. Circles: historical herbarium samples, triangles: contemporary samples. (**B**) Genetic structure estimated by pairwise *F_ST_* (weighted) between native range populations and Europe. Populations are split by time period (historical and modern). Shading of the boxes corresponds to the *F_ST_*-value, with yellow boxes indicating low *F_ST_* and purple boxes indicating high *F_ST_*.

**Fig. 2.**
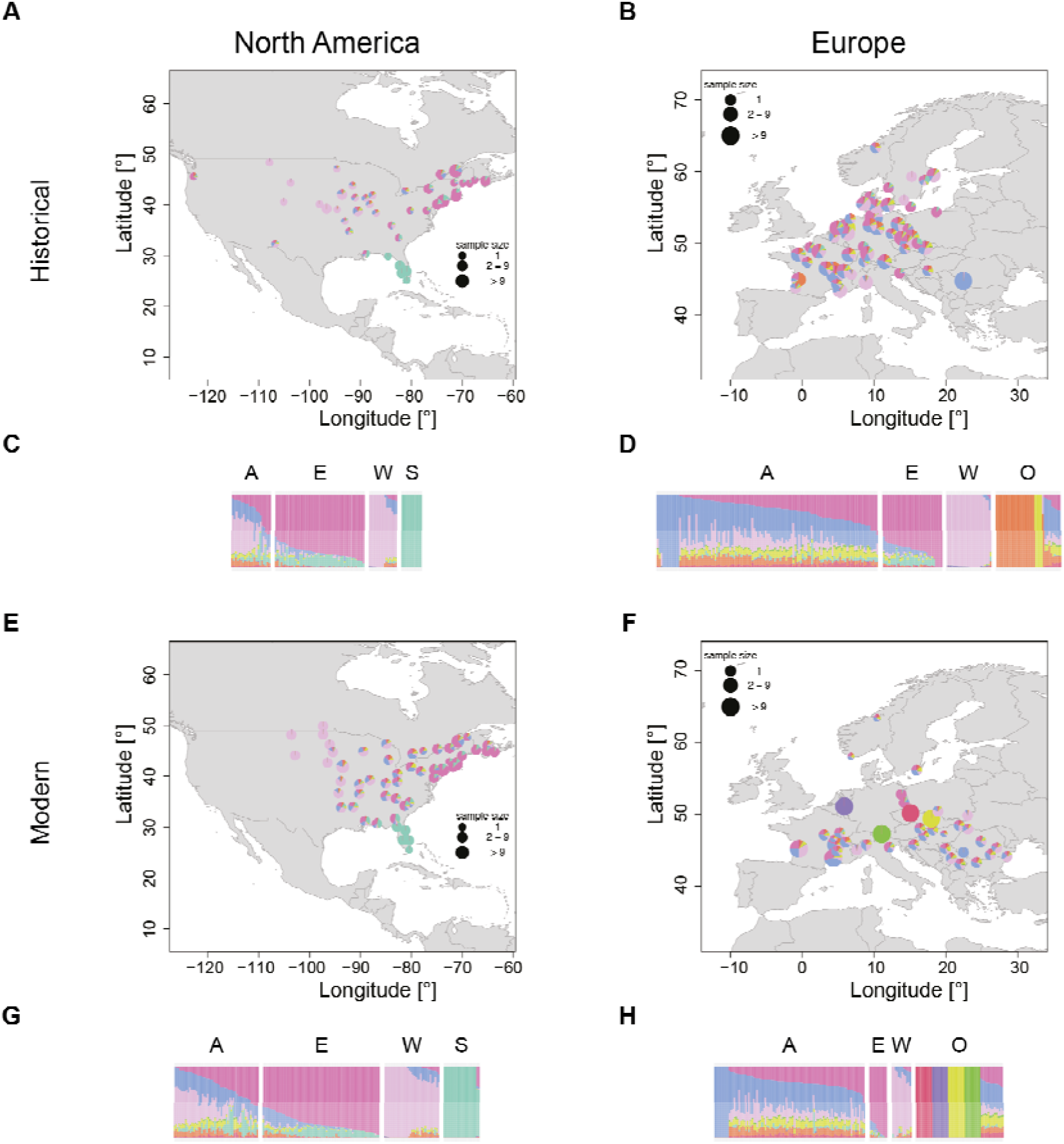
Admixture proportions for *Ambrosia artemisiifolia* populations. The NGSadmix run with the highest likelihood for *K*=9 was used for plotting and the same color scheme was used across all panels. (**A, B, E, F**) Admixture maps. Samples within 100 km were grouped together and the average ancestry across those groups was plotted. If samples were grouped together, ancestry values were plotted at the centroid of the group. (**C, D, G, H**) Admixture barplots. Each bar represents one individual. Samples are grouped based on their assignment to a genetic cluster (based on *K*=9): A: Admixed, E: East, W: West, S: South, O: other. (A) Historical North America. (B) Historical Europe. (C) Historical North America. (D) Historical Europe. (E) Modern North America. (F) Modern Europe. (G) Modern North America. (H) Modern Europe. The NGSadmix run with the highest likelihood was used for plotting and the same color scheme was used across all panels.

In the introduced European range, no clear relationships with a single native-range population are observed in both historic and modern times. European samples do not cluster with the main NA South cluster on the PCA (Fig. 1) and show overall little of the main NA South cluster in the admixture analysis. None of the European samples were assigned to the South population based on the admixture analysis for *K*=9. Furthermore, F_ST_ values are higher between Europe and NA South than between Europe and the other three North American populations. Based on F_ST_, Europe is almost equally close to the West and East population in both the historical and modern time period. Europe shows the lowest genetic distance to the Admixed population, which is lower than values between the two time periods within the same populations. In the admixture analysis, more than half of the European samples for both the historical and the modern time period are assigned to the Admixed population. The fraction of samples stays almost constant over time, with 56.7% in the historical time period and 54.7% in the modern time period. For both the East and the West population, the fraction of European samples assigned to them decreases over time. In historical times, 15.2% of samples were assigned to the East population and 11.4% to the West population. Among modern samples, only 6.5% were assigned to the East population and 7.1% to the West population. Over time, more samples within Europe could not be assigned to any North American population and several unique genetic clusters are found in modern Europe (Fig. 2). These changes are evident in the analysis of spatial groups that show drastic changes over time in the admixture analysis (Fig. 3) as well as in the analysis of pairwise F_ST_ as modern spatial groups cluster outside the North American range in the MDS analysis (Fig. 3b). Moreover, these spatial groups also cluster outside the North American range on the PCA (Fig.1). All spatial groups in Europe show signals of introgression from *A. trifida*, with the highest values found in those that form unique genetic clusters (Appeldorn, Innsbruck, Prague, Brno) (Fig. S1). In addition, two modern populations (Appeldorn and Bordeaux) show signals of introgression from *A. psilostachya* (Fig. S2). Due to the high genetic differences even of geographically close spatial groups in Europe, no isolation-by-distance pattern could be found in Europe, unlike in the native North American range (Fig. S5).

**Fig. 3.**
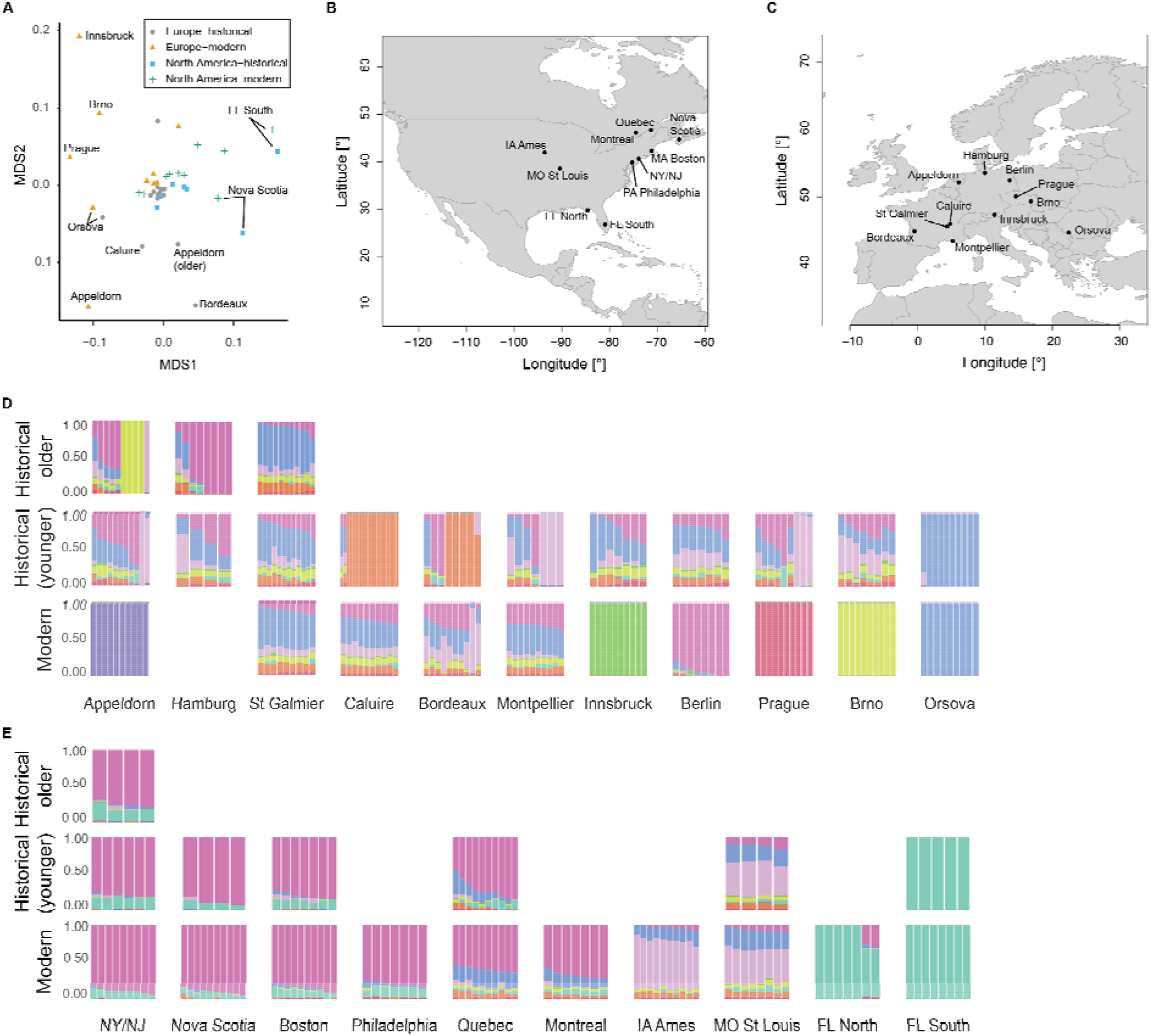
Genetic structure of spatial groups. (**A**) MDS plot of pairwise *F_ST_* between spatial groups (stress = 0.17). Gray circles: historical Europe, orange triangles: modern Europe, blue squares: historical North America, green crosses: modern North America. (**B**) Location of spat**ial** groups in North America. (**C**) Location of spatial groups in Europe. (**D**) Admixture barplots for *K*=9 of European spatial groups. (**E**) Admixture barplots for *K*=9 of North American spatial groups. (E-F) If the spatial group was split into an older (collected before 1900) and younger (collected after 1900), the top row shows the older and the middle row the younger historical time period. Otherwise, the middle row shows the historical time period. Bottom row: modern time period. For the admixture barplots, the same color scheme as in fig. 2 was used.

The highest mean heterozygosity is found in the historical Admixed population and is significantly higher (Mann–Whitney U test, *p*<0.05) than in all other populations except historical West (Fig. 4). The lowest mean heterozygosity is found in the modern Europe population and is significantly lower (*p*<0.05) than in all other populations except historical South and modern East. The lowest heterozygosity in the native range is found in modern East, which significantly differs (p-value<0.05) from all other native range populations except historical South. The effective population size (N_e_) is higher in Europe than in any of the North American populations (Table S1) and decreases over time. In the native range, the South population has the lowest and the Admixed population the highest N_e_. In the native range, N_e_ increases over time for all but the East population. Tajima’s D is negative in all populations, with the lowest value found in historical Europe, followed by historical East and historical Admixed (Table S1), and is generally lower in historical populations than in modern ones.

**Fig. 4.**
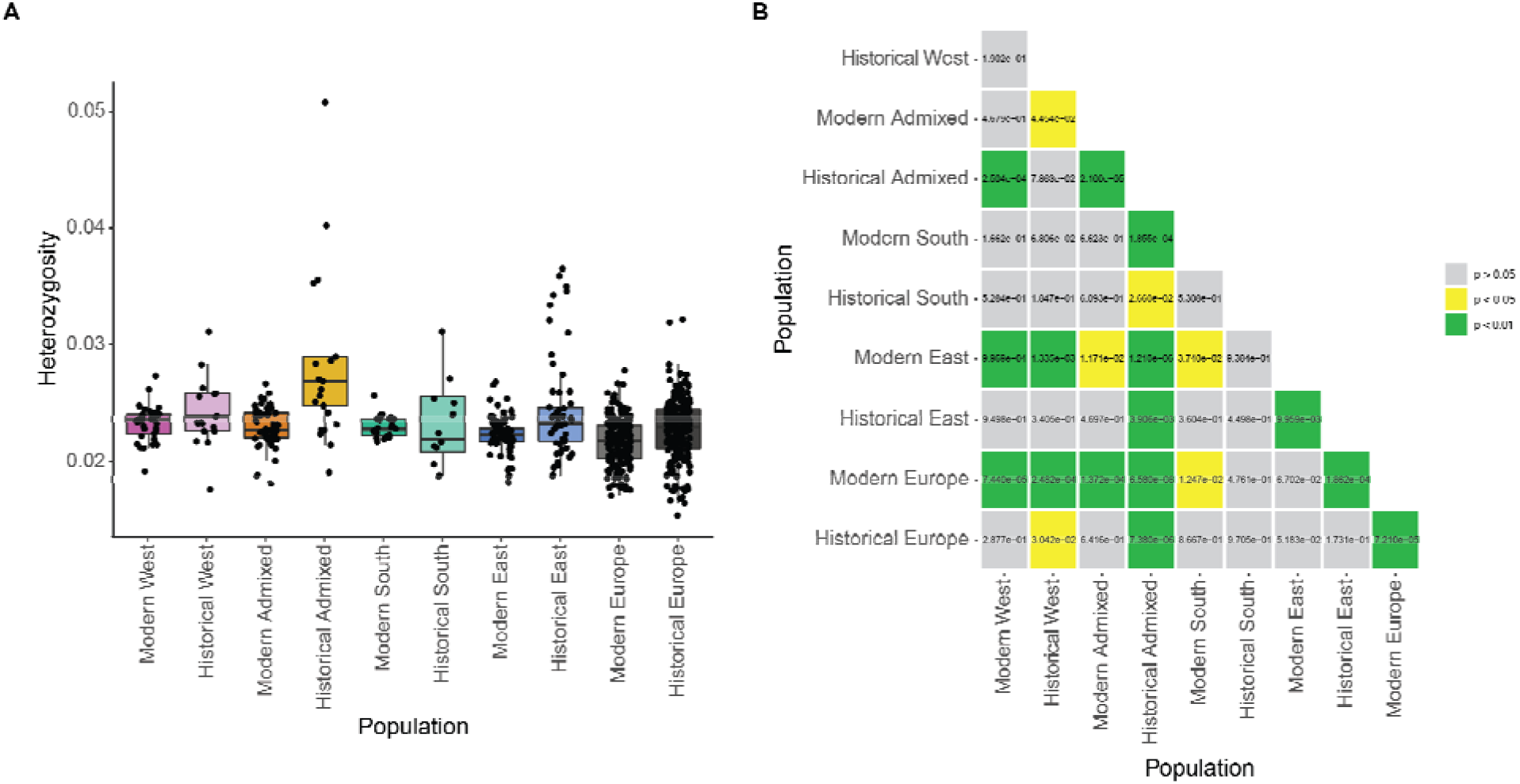
Heterozygosity. (**A**) Heterozygosity of individuals within North American and Europea**n** populations. North American populations are defined based on genetic clustering and geograph**y.** Color scheme is the same as in the PCA (Fig. 1). Samples with nuclear genome coverage below 0.5X after MAPQ 25 filtering were removed as downsampling experiments showed that the heterozygosity estimate at such low coverage diverges from the true heterozygosity (see SI). (**B**) *P-values* of Mann–Whitney U test between heterozygosity levels of different populations. Samples with coverage below 0.5X are removed*. p-values* below 0.05 are highlighted in yellow, *p-values* below 0.01 are highlighted in green.

### Selection scanning

Between historical Europe and modern Europe, a total of 353 F_ST_-outlier windows were identified. These contained 111 unique *A. artemisiifolia* genes, of which six matched flowering time genes of *Arabidopsis thaliana (43*). Between modern North America and modern Europe, a total of 442 F_ST_-outlier windows were found. These contained 139 unique genes, of which seven were homologues of *A. thaliana* flowering time genes (*43*). Of the outlier windows, 159 are shared between the comparisons of historical/modern Europe and modern Europe/North America. For both comparisons, outlier windows have on average a negative Fay & Wu’s H, which is significantly lower than in non-outlier windows (*p*<2.2e-16) while outlier windows in historical Europe and modern North America show a positive Fay & Hu’s H on average (Fig. 5). The GO enrichment analysis shows 45 enriched GO terms in the comparison of historical and modern Europe (Fig. 5, Data S1). Among other functions, these GO terms are associated with growth, stress response, light response, circadian regulation, response to phosphate and nitrate, flowering, and pollen recognition. Of these, 17 are also shared in the comparison of modern Europe and North America. Between modern Europe and modern North America, 47 GO terms are enriched in *F_ST_*-outlier windows (Fig. 5, Data S1). These include functions associated with growth, defense, response to salinity, flowering, and response to phosphate. Of the top outlier SNPs (Z>100) between historical and modern Europe, 11 out of 199 are found within eight gene regions (Data S2). Several of these genes are orthologs of well characterized genes in *A. thaliana*. Functional analysis of AT1G47740 (PPPDE putative thiol peptidase family protein) in *A. thaliana* showed that mutants are involved in abiotic stress response evidenced by expression changes in response to cold (downregulation), oxidation (downregulation) and osmotic stress (upregulation) (*44*). Mutants of AT2G13540 (*ABA HYPERSENSITIVE 1, ABH1*) are early flowering (*45*), drought resistant (*46, 47*), and hypersensitive to the plant hormone abscisic acid, which regulates development and stress response (*46*). Loss of *ABH1* function in *A. thaliana* results in abnormal processing of mRNAs for the important floral regulators *FLC, CO*, and *FLM(45*). AT5G47910 (*RESPIRATORY BURST OXIDASE HOMOLOGUE D, RBOHD*) is involved in defense response to abiotic stress and pathogens (*48, 49*), specifically via its interaction with the *AtrbohF* gene, which allows tuning the spatial control of the production of reactive oxygen intermediates and hypersensitive response around sites of infection (*49*). The protein encoded by AT3G19540 (*BOUNDARY OF ROP DOMAIN4, BDR4*) recognizes acetylation codes of histones also during mitosis and thus likely contributes to the transcriptional memory transmission to the next cell generation (*50*).

**Fig. 5.**
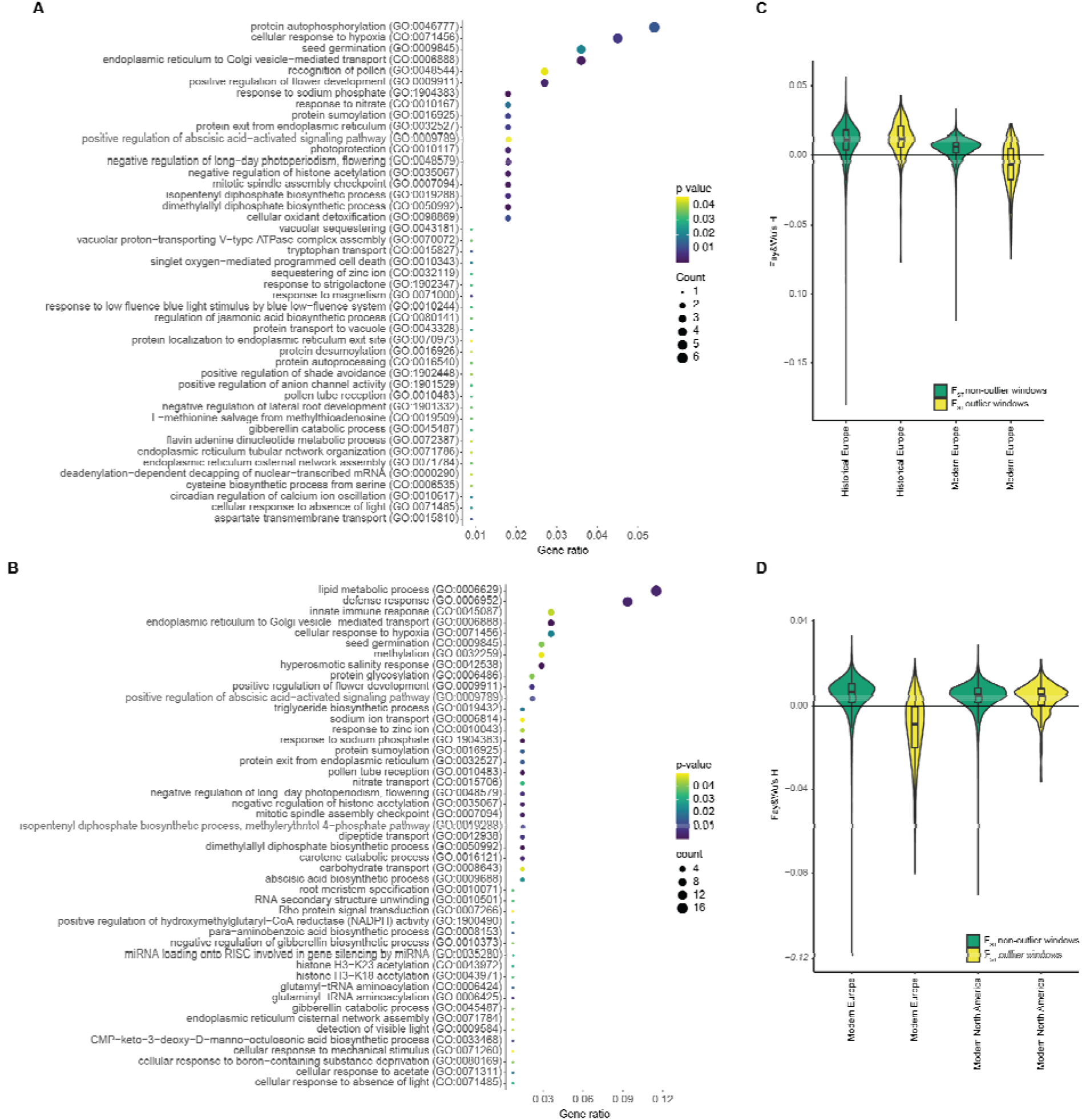
Selection scan. (**A-B**) Significantly enriched GO terms in the *F_ST_* outlier windows. The size of the circles represents the number of significant genes annotated for the respective GO term. The color represents the *p-value* for the enrichment with yellow representing high and purple low *p-values*. The gene ratio is the number of significant genes divided by the total number of genes in *F_ST_* outlier windows. (A) Enriched GO terms for historical vs. modern Europe. (B) Enriched GO terms for modern Europe vs. modern North America. (**C-D**) Fay & Hu’s H in *F_ST_* outlier (yellow) and non-outlier (green) windows. The boxplots show the median, first and third quantile. (C) Outlier windows for *F_ST_* between historical and modern Europe. (D) Outlier windows for *F_ST_* between modern Europe and modern North America.

*BDR4* is also implicated in development or function of vessels and pit boundaries in the xylem, which transports water and nutrients from the plant-soil interface to stems and leaves (*51*). The gene AT3G21670 (*NRT1-PTR FAMILY 6.4/NPF6.4*) is involved in nitrate transport (*52*).

### Pathogen identification

A total of 68 different pathogens were identified in the entire dataset (Fig. 6, Data S3). Fewer pathogens were identified in the historical herbarium samples (38 in Europe and 32 in North America) compared to the contemporary samples (58 in Europe and 60 in North America). As this difference could result from the shorter fragment length and ancient DNA damage in the historical samples, comparisons were only done within time periods. In the historical time period, eight pathogen species detected in North America are absent in Europe, while 14 pathogens detected in Europe are absent in North America. Five species have a significantly higher (*p*<0.05) prevalence in historic Europe, while one is significantly higher (*p*<0.05) in historical North America. In the contemporary samples, nine pathogens are absent from Europe and present in North America. Of these, two show the same pattern in the historical time period. Seven pathogens present in modern Europe are absent in modern North America. Of these, three show the same pattern in the historical specimens. In the contemporary time period, eight pathogens show a significantly higher (*p*<0.05) prevalence in Europe and 12 in North America. In general, either more or a significantly higher prevalence of *Xanthomonas* (Fig. 6) and *Pseudomonas* taxa are found in North America and either more or a significantly higher prevalence of *Dickeya* and *Brenneria* species are found in Europe.

**Fig. 6.**
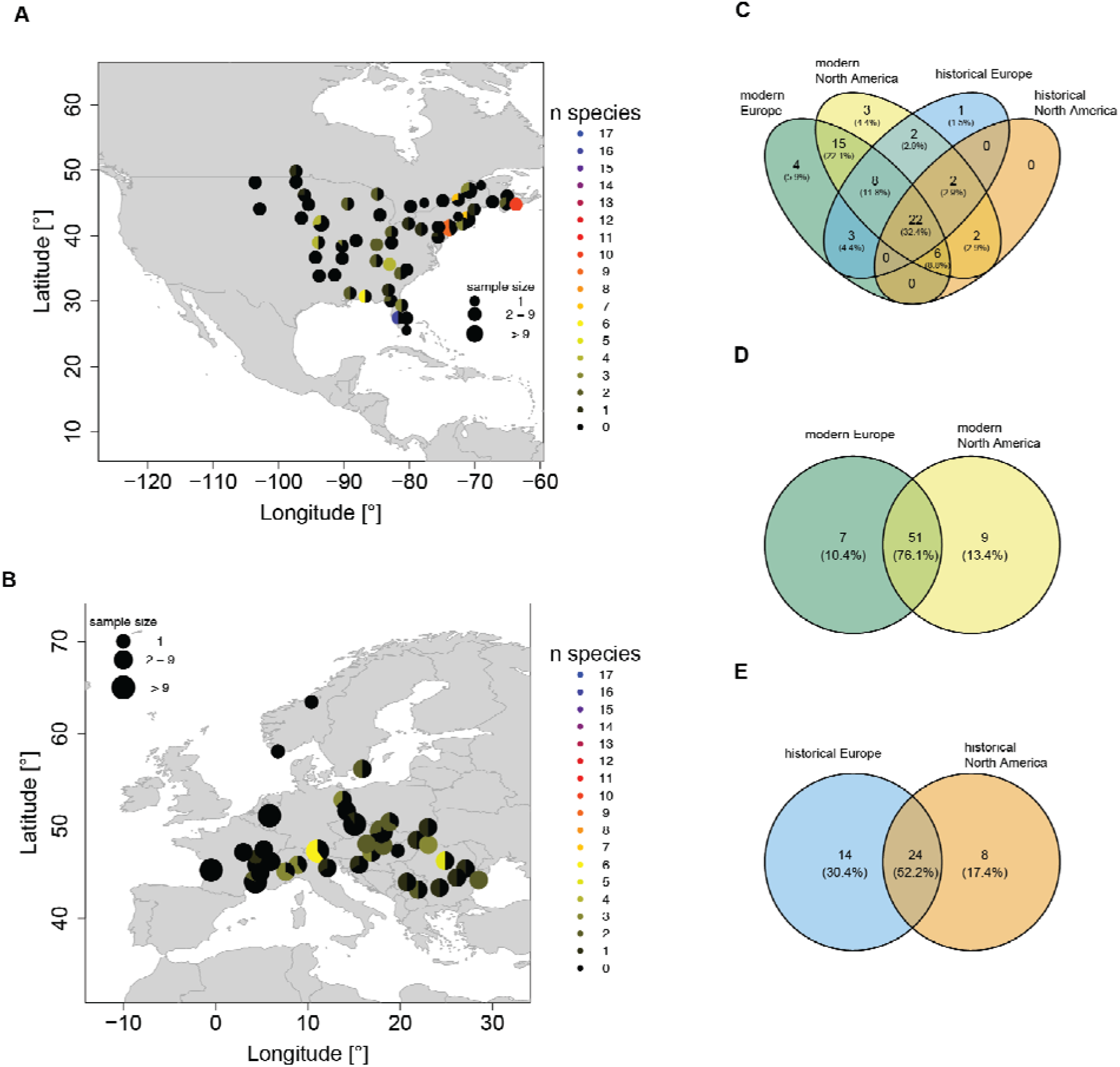
Pathogen presence. (**A-B**) Prevalence of *Xanthomonas* spp. in contemporary samples. Samples within 100 km were grouped together. The pie chart indicates the fraction of samples **in** which *Xanthomonas* spp. are present with black indicating no *Xanthomona*s species identified. The color indicates how many different *Xanthomonas* species were identified at a location. (**A**) modern North America. (**B**) modern Europe. (**C**) Venn Diagram of pathogens identified in modern European (green), modern North American (green), historical European (blue) and historical North American (orange) samples. (**D**) Venn diagram of pathogens identified in modern European (green) and modern North American (yellow) samples. (**E**) Venn diagram of pathogens identified in historical European (blue) and historical North American (orange) samples.

## Discussion

Using the largest collection of conspecific genomes derived from herbarium specimens of any species, we discovered a remarkable turnover in the genetic structure of introduced *A. artemisiifolia* populations over the brief window of time during which this plant has established itself in Europe. This finding likely reflects multiple introductions from diverse sources, drift during introduction bottlenecks, and even introgression from related species. Moreover, we found evidence of rapid adaptation and support for the role of microbial enemy release in the success of this invasive weed.

Using high-resolution spatio-temporal sampling of *A. artemisiifolia* populations in both North America and Europe, we found genomic signals of divergent selection during range expansion, pointing to rapid adaptation during the invasion. Many of these associated genes were involved in responses to stress and light, as well as flowering, defense, and growth. Our results provide a genomic foundation for understanding previous work describing major phenotypic differences between samples from Europe and the native ranges, with European populations characterized by reduced drought-resistance and a higher allocation of resources towards growth and reproduction (*37*), well in line with earlier common garden experiments that showed strong evidence of adaptation in the European range in traits such as plant size, reproduction investment, sex allocation, phenology, dichogamy, specific leaf area, and plant growth (*53*). Adaptive processes may have caused differences in early life cycle stages in European and North American populations. Germination rate, germination speed, frost tolerance of seedlings, as well as the temperature niche width for germination were significantly higher and broader for European populations (*54*). Consistent with our finding of rapid adaptation of flowering genes, European populations already show clines in flowering time, similar to those found in the native range, which likely reflect local adaptation (*39*).

It has long been suggested that escape from natural enemies in the native range can facilitate the invasion success of introduced species (*12–14*). These enemies can be animals like the ragweed leaf beetle (*Ophraella communa*), but also any of the multitudes of microbial pathogens known to affect *A. artemisiifolia* (*55*). In addition to some plant pathogen bacteria being absent or of lower prevalence in Europe (e.g. *Xanthomonas*), we also detected some taxa only in Europe or at higher prevalence than in North America (e.g. *Brenneria, Dickeya*). Bacterial plant pathogens often have high host-specificity, but can sometimes live on non-host plants without causing disease (*56*). It is thus important to ascertain whether certain pathogens actually cause disease in the plant in question before drawing conclusions about enemy-escape. *Brenneria, Dickeya*, and *Xanthomonas* are the three genera that show differences in prevalence or presence between Europe and North America. Of these, only *Xanthomonas* spp. are known to cause disease in *A. artemisiifolia*, inducing 60% mortality in infected plants (*56*), and show reduced prevalence in Europe. Dovetailing with our detection of the strongest selection signatures on the gene
*RBOHD*, a gene associated with defense against pathogens, these findings are in favour of the enemy release hypothesis and suggest that together with the release from herbivores such as the ragweed leaf beetle, escape from native-range microbes facilitated the success of *A. artemisiifolia* in Europe.

The population structure we found in the native range reenforces previous results based on reduced-representation genomic data from contemporary samples (*57, 58*). The most differentiated population is the South population in North America, which is restricted to the southeastern United States (Florida, coastal Mississippi, and Georgia). Based on the palynological records from sediment cores (*59, 60*), this region was a refugium during the last glacial maximum. If the East and West populations originate from different glacial refugia, this could explain the higher divergence of the South population from other North American populations. The South population has the lowest N_e_ in both historical and modern times. Together with the negative Tajima’s D and admixture results, this indicates that the South population experienced a population expansion but stayed relatively isolated from the other North American populations. Based on the global F_ST_, PCA and admixture results, this population did not contribute substantially to the Admixed population in the native range or to the introduced European range.

For North America, in contrast to Europe, we found clear stasis in the population genetic structure. The only exception to this was a shift in the extent of the Admixed cluster. Based on ABC-RF simulations and reduced-representation genomic data from present-day populations, van Boheemen *et al*. (*57*) estimated the Admixed population formed more than 200 years ago, thus suggesting it predated the introduction of the species to Europe. Anthropogenic disturbances, such as forest clearance and the expansion of agriculture (*29*) are the most likely cause for the formation of this Admixed population (*29*). By directly sampling historical herbarium samples, we confirm that this cluster already existed in the late 19th century, with the oldest sample in our study from this population dated to 1875. High heterozygosity values and a relatively high N_e_ for the historical Admixed population, especially compared to the modern Admixed population, indicate that the admixture event happened shortly before the period in which the majority of our historical samples were collected.

Despite being present in French botanical gardens as early as 1763, wild populations of ragweed were not reported before the late 19th century in France (*28*), and even later in other parts of Europe (*61*). Historical records and herbarium data suggest that there were several independent introductions of *A. artemisiifolia* into Europe, rather than a single introduction and a subsequent spread (*28*). These introductions likely arose from different source populations, as we find that many historical samples from Europe are fully assigned to a native-range population (11% West; 15% East; 57% Admixed). Some samples (17%) could not be assigned to any of the native genetic clusters, and the fraction of such samples increases over time to 32%. This suggests that *A. artemisiifolia* was already present in Europe well before most of our historical samples were collected, and that genetic drift associated with initially small population sizes, combined with strong selection pressure, may have led to the rapid formation of unique genetic clusters early in Europe.

It is also clear that introgression from related plant taxa contributed to the formation of these unique genetic clusters in Europe. *A. artemisiifolia* can produce hybrids with *A. trifida* (*62*) and *A. psilostachya* (*63*), both of which were introduced to Europe around the same time as *A. artemisiifolia* (*64*) and had opportunities to hybridize with *A. artemisiifolia* in the introduced range, although hybrids have not yet been reported to produce viable seeds (*31*). We find signals of introgression from both *A. trifida* and *A. psilostachya* in Europe with high levels in those populations that form unique clusters in Europe. Hybridization in the introduced range may mitigate Allee effects, which tend to be particularly strong in self-incompatible species like *A. artemisiifolia*. In addition to ‘demographic rescue’, interspecific hybridization early in the invasion may have offered *A. artemisiifolia* populations other benefits, including heterosis and the adaptive introgression of beneficial alleles. As we also find introgression in some native range populations, it is possible that the observed pattern in Europe is due to older introgression in the source population.

The origin of the European *A. artemisiifolia* invasion has been debated in the literature. All studies have found evidence for multiple introductions (*28, 36, 65*). However, some studies have suggested that the admixture was largely sourced from the native range admixed region (*29*), while others suggested the admixture occurred post-invasion (*65, 66*). Our temporally stratified snapshots of population structure provide clearer insights into this debate.

As a whole, the European population is most closely related to the North American Admixed population. Moreover, most of the European samples cluster with the Admixed population in the PCA, and more than half of the samples are assigned to the Admixed cluster in the admixture analysis. None of the samples collected before 1892 in Europe (*n*=78) were assigned to the West cluster. According to herbarium data and bibliography, the first introductions to Europe occurred over a short time period of about ten years from Eastern North America (*67*). We thus find it unlikely that substantial admixture between individuals that originated from the native-range East and West genetic cluster in the introduced range led to the observed pattern and conclude instead that the native-range Admixed cluster was the main source of introduction in Europe. The East and West populations also seem to have contributed to Europe, as several European samples show nearly complete assignment to the main West or main East genetic cluster in the admixture analysis and also group with the West or East population on the PCA. Interestingly, the prevalence of West and East cluster ancestry decreases over time in Europe, while the fraction of samples being assigned to the Admixed cluster does not. Previously it has been suggested that the population genetic differentiation between eastern and western European *A. artemisiifolia* reflects historical introduction and trade routes (*68*). In contrast, our results indicate that the main source of introduction in both western and eastern Europe is likely the Admixed population from the native range. Indeed, the oldest samples in both eastern and western Europe are frequently (57%) placed in the Admixed cluster, supporting the hypothesis that substantial admixture either occurred prior to introduction or very early in the invasion process. Due to the admixture of several native populations, much of the native-range genetic variance was introduced to historical Europe, as evidenced by no significant decrease (p>0.05) in heterozygosity levels in Europe compared to the native range and higher N_e_ than in the native range. Over time, the European population has diverged from the North American population, as fewer samples are assigned to the native genetic clusters and more genetic clusters unique to Europe have emerged.

The increasing pace of biological introduction to novel ranges via global trade and climate change severely threatens global biodiversity. Using the weed *A. artemisiifolia* as a model for plant invasion, we demonstrate that combining population genomic analysis with a metagenomic approach can identify factors that may facilitate the success of plant invaders both prior to and following the introduction event. These factors include pre-introduction admixture of different source populations in the native range, rapid adaptation, introgression from other species and the escape from some plant pathogens. We show that microbial pathogens both old and new play a role in the adaptive landscape of highly successful invasive plants like *A. artemisiifolia*. The identification of *A. artemisiifolia*’s lost native-range pathogens informs future efforts to devise effective biological control measures.

Our study illustrates the potential of global herbarium collections as a rich source of historical material for high-resolution population genomic and metagenomic investigations over continental and even global spatial scales. These meticulously curated and often well-preserved plant specimens contain not only the host plant genome, but also a complex community of associated microbes that, when considered together in a hologenomic framework, can reveal a rich history of co-evolution and networks of synergy and antagonism during the Anthropocene.

## Supporting information

DataS2

DataS3

DataS4

DataS5

## Acknowledgments

Some sequencing was performed by the NTNU Genomics Core Facility (GCF), which is funded by the Faculty of Medicine and Health Sciences, Norwegian University of Science and Technology (NTNU), and the Central Norway Regional Health Authority. Some analyses were performed on resources provided by the National Infrastructure for High Performance Computing and Data Storage in Norway (UNINETT Sigma2). The authors acknowledge support from Science for Life Laboratory, the Knut and Alice Wallenberg Foundation, the National Genomics Infrastructure funded by the Swedish Research Council, and the Uppsala Multidisciplinary Center for Advanced Computational Science for assistance with massively parallel sequencing and access to the UPPMAX computational infrastructure. We thank the curators from the following herbaria for allowing us to destructively sample their collections: B, BR, BRNU, C, FI, G, GH, GOET, GZU, HBG, I, IASI, JE, L, LD, LY, MARS, MASS, MO, MPU, NEBC, NEU, NY, P, PH, PR, PRA, PRC, QFA, S, STU, TRH, UPS, US, W, WU. We thank Andrew Foote for useful discussions about data analysis.

## Funding

NTNU Onsager Fellowship award (MDM)

Norwegian Research Council Young Research Talents award 287327 (MDM)

SYNTHESYS Project (www.synthesys.info, financed by European Community Research Infrastructure Action under the FP7 “Capacities” Program) (MDM)

Monash University startup grant (KH)

Australian Research Council Discovery Project award DP180102531 (KH)

Human Frontiers Program Grant RGP0001/2019 (KH)

Natural Sciences and Engineering Research Council of Canada (NSERC) grant #353026 (LHR)

the Luomus Trigger Fund (Finland) (PP)

Systematics Research Fund (UK) (PP)

## Author contributions

Conceptualization: MDM, KH

Sample Collection: VCB, MDM, KH, FB, LvB, MG, HMS, SL, GK, BC, YS, PP

Resources: MDM, KH, JCB, SG, LD, LHR, MTPG

Methodology: VCB, CL, KN, GLO, LvB, JYL, FLK, PP, MDM

Investigation: VCB, PB, BP, XS, JW

Visualization: VCB

Funding acquisition: MDM, KH, LHR, PP

Project administration: MDM, KH

Supervision: MDM, KH

Writing – original draft: VCB, MDM, KH, with input from all co-authors

Writing – review & editing: all co-authors

## Competing interests

Authors declare that they have no competing interests.

## Data and materials availability

DNA sequences generated for this study can be found under ENA study PRJEB48563. Previously published data can be found under ENA studies PRJNA339123 and PRJEB34825. A complete list of accession codes for each sample can be found in table S5. The programs, *R* packages and functions used are described in detail in the Methods with citations. The scripts and commands that call these tools are available upon request.

## Supplementary Materials

Materials and Methods

Figs. S1 to S21

Tables S1

References (*69–118*)

Data S1 to S5

## Notes

### Competing Interest Statement

The authors have declared no competing interest.

