## Supplementary Information for "Uncovering the hologenomic basis of an extraordinary plant invasion"

### Hologenomic analysis of hundreds of herbarium specimens reveals the genomic basis of plant invasion

**This PDF file includes:**

Materials and Methods
Figs. S1 to S21
Table S1
Captions for Data S1 to S5

**Other Supplementary Materials for this manuscript include the following:**

Data S1. Significantly enriched GO terms in Fst-outlier windows.
Data S2. Top outlier SNPs (Z>100) located in gene regions.
Data S3. Prevalence of plant pathogens in the different ranges and time periods.
Data S4. Overview of herbarium and contemporary *Ambrosia artemisiifolia* samples.
Data S5. Overview of outgroup samples from different *Ambrosia* species.

### Materials and Methods

*De-novo* assembly of reference genome

Germinated seeds were collected from a natural *Ambrosia artemisiifolia* population in North Dakota, USA (46.298° latitude, -103.918° longitude) in 2008, the individual plant AA19_3_7 was grown to maturity in a greenhouse at the Department of Botany, University of British Columbia. Fresh leaf tissue was sampled, flash-frozen in liquid nitrogen and then stored at -80°C. High-molecular-weight (HMW) genomic DNA was extracted using a CTAB protocol (*69*, *70*) and prepared for short-insert sequencing and mate-pair sequencing. Sequencing was performed at Genome Quebec, generating four lanes of short-insert data from two TruSeq libraries and two lanes of Nextera mate-pair data from two libraries (5-kbp and 10-kbp inserts) on an Illumina platform. After adapter trimming with Trimmomatic v0.38 (*71*), the data was assembled using *Meraculous-2D* v2.2.6 (*40*) with the command-line options *diploid_mode 2*, *no_strict_haplotypes 0*, *genome_size 2*, *gap_close_aggressive 1*, *mer_size 55*, and *min_depth_cutoff 11*.

*Ambrosia artemisiifolia* is self-incompatible, so it was not possible to obtain a homozygous plant for sequencing and assembly of a reference genome. Due to the heterozygous nature of the genome, our initial assembly contained many near-duplicate haploid contigs (haplotigs) from individual AA19_3_7's homologous chromosomes. We attempted to remove these haplotigs from the reference assembly to ensure the length of the reference genome is not falsely inflated using the software *PurgeHaplotigs* (*72*). After producing a histogram of contig read depth, low, mid-point and high cutoffs were designated as read depth values of 20, 110 and 195 respectively. Alongside a junk threshold of 80% (-j 80) and a suspected haplotig threshold of 60% (-*s 60*), these values were used to create a pool of suspect haplotigs. A final step was performed to split duplicate contigs into a haplotig pool using a threshold of 70% (-*a 70*) identity to confirm a contig as a haplotig, and then to create a new, filtered assembly with the haplotigs removed. Comparison of basic single copy orthologous (BUSCO) genes revealed a large increase in single copy genes after purging, using *BUSCO* v5.1(*42*). Final BUSCO scores after purging were 237 complete buscos (136 single copy, 101 duplicated, 15 fragmented and 3 missing) out of 255 BUSCO groups searched. This was provided as the input assembly for the genome assembly scaffolding described below.

Additional HMW genomic DNA was provided to the commercial provider Dovetail Genomics, which prepared two Chicago libraries as described previously (*73*). Briefly, for each library, ~500 ng of HMW gDNA was reconstituted into chromatin in vitro and fixed with formaldehyde. Fixed chromatin was digested with MboI, the 5' overhangs filled in with biotinylated nucleotides, and then free blunt ends were ligated. After ligation, crosslinks were reversed, and the DNA purified from protein. Purified DNA was treated to remove biotin that was not internal to ligated fragments. The DNA was then sheared to ~350 bp mean fragment size and sequencing libraries were generated using NEBNext Ultra enzymes and Illumina-compatible adapters. Biotin-containing fragments were isolated using streptavidin beads before PCR enrichment of each library. The Chicago libraries were then sequenced on an Illumina HiSeq 2500 platform. The number and length of read pairs produced for each library was: 256 million, 2x101 bp for library 1; 103 million, 2x101bp for library 2. Together, these Chicago library reads provided 25.6X physical coverage of the genome (1-100 kbp pairs).

The input de-novo assembly and Chicago library reads were used as input data for HiRise, a software pipeline designed specifically for using proximity ligation data to scaffold genome assemblies (*73*). Chicago library sequences were aligned to the draft input assembly using a modified SNAP read mapper (http://snap.cs.berkeley.edu). The separations of Chicago read pairs mapped within draft scaffolds were analyzed by HiRise to produce a likelihood model for genomic distance between read pairs, and the model was used to identify and break putative misjoins, to score prospective joins, and make joins above a threshold. After scaffolding, shotgun sequences were used to close gaps between contigs.

Annotation of reference genome

From sample AA19_3_7 we harvested each of four tissue types (leaf, stem, male flowers and root) by placing the tissue into aluminum envelopes and then into liquid nitrogen. The frozen tissue samples were ground using a mortar and pestle and total RNA purified using commercially prepared TRIzol reagent. The total RNA extracts were visualized using a Bioanalyzer instrument using a total RNA 6000 chip. The Bioanalyzer generates an RNA Integrity Number (RIN) that assesses the integrity of the total RNA sample, and total RNA extracts passed quality control if the RIN was $\geq 7.5$. The total RNA extracts from each tissue type were sent to the Genome Quebec Innovation Centre at McGill University where each one was prepared into a barcoded and stranded TruSeq mRNA library with an insert size of 165 bp (+/- 10%). The four completed libraries were pooled in equal amounts and sequenced on one lane of an Illumina HiSeq 2000 instrument. The RNAseq data was first cleaned using Fastx (Hannon Lab), and then a de-novo reference was assembled with all four libraries using Trinity (*74*).

Repeats and low-complexity DNA sequences were masked in the genome before gene annotation using *RepeatMasker* version 4.1.0 (*75*) using the species repeat database 'Asteraceae' with *Repbase* database version 20170127. Remaining specific repetitive elements were predicted de-novo using *RepeatModeler* version 2.0.1(*76*) on the masked genome. Subsequently, a second round of *RepeatMasker* was run with the model generated from *RepeatModeler* as custom library input on the previously masked genome. Genome annotation was performed using the genome annotation pipeline *MAKER2* version 2.31.9 (*77*) with ab-initio and homology-based gene predictions. 3,819 unique protein sequences from asterids (a monophyletic group of flowering plants), Asteraceae (sunflower family), and Ambrosia (ragweeds) were used for homology-based gene prediction. As no training gene models were available for *A. artemisiifolia*, we used *CEGMA* (*78*) to train the ab-initio gene predictor *SNAP* (*79*). *MAKER2* was run with command-line arguments *model_org=simple*, *softmask=1*, *augustus_species=arabidopsis* and the *snaphmm* parameter was set to the HMM generated in the manual training of *SNAP*. As expressed sequence tag (EST) evidence, we used the *Trinity*-assembled transcriptome. The 34,066 predicted proteins were then compared with *Arabidopsis thaliana* annotations (TAIR10 representative gene model proteins (*80*)) using the *blastp* command in *BLAST+* (*81*). 29,927 predicted *A. artemisiifolia* proteins matched *A. thaliana* genes with an E-value < 1×10-6; these annotations were retained for downstream analysis. Additionally, annotations were cross-referenced with 306 *A. thaliana* flowering time genes (*43*). 566 predicted *A. artemisiifolia* genes were matched to this dataset, representing 191 unique *A. thaliana* flowering time genes.

Acquisition of contemporary samples

Silica-dried leaf tissue samples were obtained from wild populations in North America and Europe via the authors' personal collections and a network of collaborators. Some samples were provided as seeds, which were sown and raised in the greenhouse facilities at the NTNU University Museum's Ringve Botanical Garden (Trondheim, Norway). From these plants, leaf tissue was harvested and silica dried. See supplementary table of sample provenance (Data S4).

Acquisition of historical herbarium samples

To summarize and select from available *A. artemisiifolia* herbarium specimens, we made a list of samples already available from previous studies (*29,37,57*), those we found in online databases of herbarium collections (e.g. the Global Biodiversity Information Facility, GBIF, www.gbif.org) and by directly contacting herbaria. We preferred specimens that were collected before 1940 to represent the initially introduced populations into Europe. When the collection latitude and longitude coordinates were not available as metadata, this information was inferred from the centroid (as defined by the web tool Google Maps) of the most specific geographic sampling location (e.g. the city) described on the herbarium voucher sheet. A complete list of samples included in this study can be found in Data S4, along with sample locations and assignment to broader populations (as described below). To evaluate if specimens in our dataset were misidentified and to test for introgression of some species, we included shotgun-sequencing data of 123 samples of other *Ambrosia* species (hereinafter referred to as outgroup samples), including samples of known hybrids of *A. artemisiifolia* with other *Ambrosia* species (Data S5).

DNA extraction, library preparation and sequencing of herbarium samples

For this study, we generated new shotgun sequencing data for 382 historical herbarium and 366 contemporary specimens of *A. artemisiifolia* and combined it with already published data (*82,83*). 77 herbarium samples and 16 contemporary samples were later removed for various reasons (see sections below), leading to a final dataset of 655 samples. A full overview of sample sources, laboratory methods and inclusion in the final dataset can be found in Data S4. Sampling locations are displayed in Fig. S6.

For the historical herbarium samples processed for this study, all pre-PCR steps were carried out in a dedicated, positively pressurized ancient DNA laboratory at the NTNU University Museum. DNA was extracted from leaf tissue using the Qiagen DNeasy Plant Mini Kit according to the manufacturer's instructions except for the addition of an overnight incubation step with proteinase K as previously described (*29*). DNA concentration was quantified with a Qubit 2.0 fluorometer using the BR dsDNA kit.

DNA extracts were converted into blunt-end double-stranded Illumina libraries using the BEST protocol (*84*), in which custom blunt-end adapters (*85*) were ligated to the DNA fragments, or single-stranded Illumina libraries using the Santa Cruz Reaction (SCR) protocol (*86*). During indexing PCR, custom index primers were used to generate dual-index libraries. Indexing PCR was carried out either in a 50 µL or a 100 µL reaction with 5-10 µL library template, 0.2 mM each dNTP, 0.2 µM sample specific forward index primer, 0.2 µM sample specific reverse index primer, 0.05 U/µL AmpliTaq Gold DNA polymerase, 1X AmpliTaq Gold buffer, 2.5 mM MgCl2, 0.4 mg/mL BSA and the rest of the reaction volume filled up with molecular grade water. The PCR was performed with an initial denaturation of 10 min at 95 ºC, then X cycles of 30 s denature at 95 ºC, 1 min annealing at 60 ºC, 45 s extension at 72 ºC, followed by a final extension of 5 min at 72 ºC. The optimal number of PCR cycles X was selected for each sample based on qPCR. Amplified libraries were purified with SPRI beads (*87*) and eluted in 33 µL Qiagen EB buffer. For some samples, two indexing PCRs were performed to increase library complexity. Samples were pooled and sequenced on Illumina platforms (see table S1 for details about the sequencing platforms used).

DNA extraction, library preparation and sequencing of modern samples

For contemporary samples, leaf tissue was collected and stored in silica gel desiccants (*88*) at room temperature until required for DNA extraction. Approximately 20-30 mg of dried leaf tissue from each sample was placed inside a 2.0 mL tube with a 3 mm stainless steel bead, and ground with a TissueLyser II (Qiagen). The DNA was extracted using a modified CTAB protocol (*89*) adapted for a 96-well plate format (*90*) using EconoSpinTM filter plates, and the DNA was suspended in 60 µL of elution buffer. Extracted DNA was quantified using a Qubit 2.0 fluorometer (Invitrogen, Carlsbad, CA, USA) using the high sensitivity dsDNA kit.

Extracts were converted into blunt-end Illumina libraries as described above. Indexing PCR was carried out in a 100 µL reaction using 7.5 µL DNA template, 0.2 mM each dNTP, 0.2 µM sample specific forward index primer, 0.2 µM sample specific reverse index primer, 1X Herculase Fusion II DNA polymerase, 1X Herculase II reaction buffer and the remaining volume filled up with molecular grade water or in a 50 µL reaction with 5 µL library template, 0.2 µM sample specific forward index primer, 0.2 µM sample specific reverse index primer, 1X Platinum SuperFi PCR master mix and the rest of the volume filled up with molecular grade water. The PCR with Herculase was performed with an initial denaturation of 3 min at 95 ºC, followed by 12 cycles of 20 s denaturation at 95 ºC, 20 s annealing at 60 ºC, 40 s extension at 72 ºC, followed by a final extension for 5 min at 72 ºC. Amplified libraries were purified with SPRI beads (*87*) and eluted in 33 µL EB buffer. The PCR with SuperFi was performed with an initial denaturation of 3 min at 98 ºC, followed by 12 cycles of 20 s denature at 98 ºC, 1 min annealing at 60 ºC, 45 s extension at 72 ºC, followed by a final extension of 5 min at 72 ºC. Amplified libraries were purified with SPRI beads and eluted in 33 µL EBT buffer. Samples were pooled and sequenced on Illumina NovaSeq. Additionally, 44 samples were sequenced on the DNBSEQ-G400 platform. See table S5 for details about the sequencing platform used.

Sequence alignment to reference genome

Raw reads were processed with the *paleomix* v.1.2.13.8 BAM pipeline (*91*). *AdapterRemoval* v2.3.1(*92*) was used to remove sequencing adapters and reads with a minimum overlap of 11 bases were collapsed into one read and treated as single-end reads during the mapping. Sequences were aligned to the *A. artemisiifolia* reference genome assembly and against the *A. artemisiifolia* chloroplast reference genome (GenBank: MG019037.1) using *bwa* v0.7.17 *mem* (*93*, *94*) without filtering for quality. PCR duplicates were marked using either *picardtools MarkDuplicates* v2.21.2 (http://broadinstitute.github.io/picard) or *bammarkduplicates* from the *biobambam2* v2.0.87 package. *MapDamage2* (*95*) was used to calculate the frequencies of base misincorporation for historical samples. The *paleomix* summary files were used to obtain mapping statistics. The endogenous content was estimated as the fraction of raw reads mapping against the *A.artemisiifolia* reference genome. Statistical analysis was conducted in *R*. The mean sequencing depth of the nuclear genome after mapping quality filtering (MAPQ ≥ 25) was 1.4X for historical herbarium samples and 2.9X for modern samples.

Genotype likelihood estimation (nuclear genome)

Genotype likelihoods were estimated for the nuclear genome with *angsd* v0.931 (*96*) with the options *-doGLF 2*, *-SNP_pval 1e-6*, *-doMaf 3*, *-doGeno -1*, *-doPost 1*, *-minMapQ 25*, *-minQ 20*, *-trim 5*, *-minMaf 0.05*, *-geno_minDepth 2*, *-setMinDepthInd 2*, *-postCutoff 0.95*, *-remove_bads 1*, *-uniqueOnly 1*, *-doPlink 2*. Only sites with sequence data for at least half of the individuals were considered. Genotype likelihoods were first calculated on the historical herbarium dataset and the contemporary dataset separately to identify positions that were variable in both.

*CallableLoci* from *GATK* v3.7-0 (*97*) was used to estimate which parts of the nuclear reference genome were reliably mappable based on mapping quality and sequencing depth distribution as described below. As the read length of historical and contemporary samples differs significantly, *CallableLoci* was run on historical and contemporary samples separately. The BAM files of 14 historical and 12 modern samples from both the native North American and introduced European range were merged with *samtools merge* v1.6 (*98*). Samples that represent the whole range and with similar sequencing depth (around 2X) were used for merging. The sequencing depth of the merged BAM files was calculated with *samtools* v1.6 depth with filtering for base quality of 20 and mapping quality of 25. The average sequencing depth was 24.3X for the merged historical BAM file and 23.0X for the merged modern BAM file. *CallableLoci* was run with a minimum base quality of 20, a minimum mapping quality of 25, a minimum depth of one third of the average depth, and a maximum depth of two times the average depth. Regions with excessive sequencing depth and with low mapping quality were removed from the resulting bed files, the bed file for the modern and historic samples combined, and overlapping regions merged with *bedtools merge* v2.25.0.

Variant sites that were in regions with low mapping quality, as well as those within regions of excessive sequencing depth identified with *CallableLoci* as described above, were removed as these regions might originate from the mitochondrial or the chloroplast genome or are gene duplications and thus violate the assumption of a diploid site in the genotype likelihood estimation. The average sequencing depth in the remaining regions was 2.3X for historical herbarium samples and 4.2X for contemporary samples. Additionally, only sites that were variable in both the historical herbarium dataset and the contemporary dataset, based on the genotype likelihood estimation on historic and contemporary samples separately, were extracted from the beagle file of the joint genotype likelihood estimation and used in the downstream analysis unless otherwise stated.

The genotype likelihood estimation was once performed on the whole dataset, including other *Ambrosia* species (see Data S5) to identify possible hybrids and misidentifications in the dataset. Based on the PCA, a total of 50 samples were removed (see Data S4). In addition, 16 samples were removed because they were either first or second-degree relatives (based on the kinship analysis described below), 23 samples were removed because of too low sequencing depth (<0.1X after filtering for MAPQ 25), and four samples were removed because of a possible sample mix-up in the lab. The genotype likelihood estimation was repeated on the reduced dataset containing 655 samples.

PCA, kinship and admixture analysis

*PCangsd* v.0.95 (*99*) was used to generate the covariance matrix and a kinship matrix for the whole dataset. The analysis was run until convergence to a minor allele frequency tolerance of 0.0001. For the PCA, the *R* function *prcomp* was used with the covariance matrix. Based on the kinship matrix from *PCangsd*, first- and second-degree relatives were removed from the analyses. For each pair of related samples, the one with the highest mean sequencing depth was kept. To identify possible hybrids or mis-identifications, a PCA based on the nuclear genome including different *Ambrosia* species was performed. Samples that clustered outside the main *A. artemisiifolia* group (PC1 > -1, PC2 < -0.35, and PC2 > 0 & PC1 < 0) were excluded from further analyses (Fig. S7). Some outgroup samples also clustered with the main *A. artemisiifolia* cluster. This might be due to mis-identifications of outgroup samples, but is more likely the result of low sequencing depth (< 0.1 X) in these samples. In total, 47 samples (6.3% of all samples) were excluded: 34 historical European (13%), 10 historical North American (8.5%) and 3 contemporary North American (1.5%) samples (Data S4). These outgroup samples were therefore excluded in the consideration of mis-identified *A. artemisiifolia* samples. The PCA was repeated including only the final set of samples. For the admixture and PCA analyses, sites in LD ≥ 0.5 were removed. LD was estimated using *Plink* v1.90 (*100*) with a window size of 50 and a step size of 5. In addition, only sites variable in both historical and modern samples were used. A total of 1,094,260 sites were used for the PCA and admixture analysis after filtering for *MAF* (minor allele frequency) 0.05. *NGSadmix* (101) was run on the reduced dataset with up to 15 ancestral populations (*K*). Ten independent runs with different seeds were performed for each *K* value. *CLUMPAK Distruct* (*102*) was used to align the output files for different *K* values. The run with the highest likelihood for each *K* was used for plotting. *K*=9 was chosen for the main manuscript as it was the most likely according to the MAP test in *PCangsd* and showed a peak with the *delta-K* method (*103*). Plots for the other *K* values can be found in the SI (Fig. S8-S20). Admixture results correlate with geographic location of the samples and were thus used to group samples. For the grouping, admixture proportions for *K*=9 were used. Samples that had at least 55% ancestry of the dominant genetic cluster in the South, East or West of North America respectively, were assigned to the South, East or West population. Of the remaining samples that were not assigned to one of the dominant South, East or West genetic clusters, that had a combined ancestry of at least 70% of the East, West, South and a fourth genetic cluster dominant in the geographic region between the three extremes were assigned to the Admixed population. Samples from the introduced European range were also assigned to these populations in the admixture analysis but were kept as a separate population for all between-population analysis. See Fig. S6 for the population assignment.

Ancestral state estimation

To generate the ancestral state for common ragweed, shotgun sequencing reads of two closely related (*104*) species were used (*Ambrosia chamissonis* and *Ambrosia carduacea*) and mapped against the *A. artemisiifolia* reference genome. The sequencing depth after MAPQ 25 filtering for both samples is 3.5X. To generate the ancestral state *fasta* file, *angsd* v0.931 (*96*) -*doFasta 2* was used with a minimum base quality of 20, minimum mapping quality of 25 and the options -*remove_bads 1*, -*uniqueOnly 1*, and -*explode 1*.

Heterozygosity, $F_{ST}$ and $N_e$ estimation

For each sample, heterozygosity was estimated over the whole genome. First, the site allele frequency (SAF) was estimated with *angsd* v0.931 (*96*) using *-dosaf 1, -minMapQ 25, -minQ 20, -remove_bads 1, -uniqueOnly 1, and -trim 5*. The site frequency spectrum was polarized using the ancestral state estimated from *A. chamissonis* and *A. carduacea*. To test if differences in heterozygosity are due to differences in sequencing depth between modern and historical samples, all samples with sequencing depth above 1X, 0.75X, 0.5X and 0.25X respectively after filtering for mapping quality 25 were downsampled to ~1X, ~0.75X, ~0.5X and ~0.25X sequencing depth with the *-downsample* option during SAF estimation. The heterozygosity estimates including all reads for each sample and downsampled to 1X, 0.75X, and 0.5X sequencing depth are strongly correlated (Fig. S20). The correlation when downsampling to 0.2X is lower, with an $R^2 < 0.9$. Thus, samples with sequencing depths below 0.5X after MAPQ 25 filtering were removed from the analysis to avoid bias due to low sequencing depth. To test if there are significant differences in heterozygosity between populations, a Mann–Whitney U test was performed in *R* v3.4.4.

For the $F_{ST}$ estimation, samples were grouped by populations (East, West, South, Admixed, Europe; Fig. 1C and 1D) and by time (historic, modern). Population assignment was based on admixture results for $K$=9 (**see admixture analysis**). Four samples from three sampling locations were assigned to either the Admixed or East population but were geographically not located in that population (Fig. S6). These samples were excluded from their respective population for all population based analysis. Samples that are first- or second-degree relatives and those that are misidentifications or possible hybrids based on the PCA analysis including outgroups from the *Ambrosia* genus were removed. First, the SAF was estimated with *angsd* using the command line options *-minMapQ 25, -minQ 20, -remove_bads 1, -uniqueOnly 1, and -trim 5*. The site frequency spectrum was polarized using the ancestral state estimated from *A. chamissonis* and *A. carduacea*.

To estimate the effective population size $N_e$, thetas were calculated with the *angsd realSFS saf2theta* tool for each population, polarized using the ancestral state estimated from *A. chamissonis* and *A. carduacea*. The mean of the Watterson estimator across the genome was used to calculate $N_e$ and the 95% CI of the mean was used to calculate the error of $N_e$. A mutation rate of $1 \times 10^{-8}$ substitutions/site/generation (estimate from the closely related species *Helianthus annuus* (*105*)), and a generation time of one year was used, as *A. artemisiifolia* is an annual plant.

Definition of spatial groups

Samples within a radius of 100 km were clustered into spatial groups using the *R* package *geosphere* and the centromere of the radius was used as the location of the spatial groups. If a spatial group contained at least four samples, we selected that group as a "population" in which changes in the genetic structure potentially could be directly observed. For some spatial groups, at least four samples were available from before 1900 and from between 1900 and 1940. For those spatial groups, we split the historical spatial group into an older (< 1900) and a younger (1900 - 1940) spatial group for a higher temporal resolution. To create modern geographic groups, we selected from available modern populations that were collected for previous studies or collected new samples in close proximity to the historical spatial groups. For one spatial group (Hamburg), no modern samples could be found due to successful eradication of the invasive plant. In North America, we chose some additional modern locations where less than four historical samples were available. We used these locations to gain a better resolution.

Pairwise $F_{ST}$ between spatial groups was calculated with *realSFS* from *angsd* (*96*, *106*). The SFS was polarized using the ancestral state. Geographic distance between the centroid of the spatial groups was calculated with the *distm* function in *R*. An MDS analysis was performed on the $F_{ST}$ distance matrix in *R* with maxit=5000. For the isolation-by-distance (IBD) analysis, a mantel test was performed using the *gl.ibd* function of the *R* package *dartR* in *R* v4.1.1 with 999 permutations. IBD was tested within historic North America, modern North America, historic Europe, and modern Europe.

To test for introgression of *Ambrosia trifida* and *Ambrosia psilostachya*, the multipopulation D-statistic (*Abbababa2*) within *angsd* (*96*, *107*) was used. *Ambrosia carduacea* was used as an outgroup as its distribution (Western North America (*108*)) does not overlap with *A. artemisiifolia* and thus introgression from this species is unlikely. D statistics of the form (E, N, *A. trifida, A. carduacea*), (N, N, *A. trifida, A. carduacea*), (E, N, *A. psilostachya, A. carduacea*), and (N, N, *A. psilostachya, A. carduacea*) were considered, where N is a spatial group from North America and E a spatial group from Europe.

<u>TreeMix</u> <u>and</u> <u>PSMC</u>

To infer effective population size change history of ragweed, we used the Pairwise Sequentially Markovian coalescent (*PSMC*) model (*109*). The sample used for *PSMC* was QC-2-30 with an average sequencing depth of 10X. To generate the diploid consensus sequence, we applied the sequencing depth filter of > ⅓ and < 2X of the average sequencing depth and filtered reads with a minimum mapping quality of 25 and a minimum base quality of 25 using samtools (*98*). Only scaffolds longer than 100 kbp were used for *PSMC*. *PSMC* was then applied to infer the density of the time to the most recent common ancestor between the two haploids across the genome. Demographic was calculated assuming a generation time of 1 year (*31*) and mutation rate of $1.0 \times 10^{-8}$ substitutions per site per generation (*105*). The robustness of the inference was estimated with 100 bootstraps (Fig. S4).

*TreeMix* v1.13 (*110*, *111*) was used to infer the phylogenetic context of different populations and other *Ambrosia* species and infer possible gene flow between populations. We defined two sets of populations including modern and historical ragweed and selected samples for each population based on our genetic admixture result (Fig. 2, Fig S6). For each population, allele frequency was estimated using *angsd* excluding sites with excessive coverage or low mean MAPQ scores and only including sites that were covered in at least two thirds of the samples by reads with MAPQ≥25 and a minimum base quality of 20. Reads that had multiple best hits as well those tagged as not primary alignment, failure or duplicate reads were removed. The reference base was assumed to be the major allele (*-doMajorMinor 4*) to ensure the same allele is the major allele across all populations. In addition, triallelic sites were excluded. *TreeMix* was run on the two sets of populations assuming 0 to 3 migration events (*m*=0-3). For each migration event number, we ran 100 replicates with different random seeds and chose the best replicated with the highest likelihood. In order to account for linkage disequilibrium, we grouped together SNPs (*-k*) using block size of 2000 SNPs for modern dataset and 1000 SNPs for historical dataset. We turned off sample size correction (-*noss*) to reduce the number of zero length branches.

Selection scanning

To identify genes putatively under selection in Europe, $F_{ST}$ was estimated in non-overlapping sliding windows with a window size of 10 kbp between historical and modern Europe, as well as between modern Europe and modern North America. As the South population did not seem to have contributed to the European invasion, it was excluded from the North American population for this analysis. The site frequency spectrum for each population was estimated with *angsd* (*96*, *106*), excluding sites with excessive coverage or low MAPQ and only including sites that were covered in at least two thirds of the samples, with MAPQ≥25, and a minimum base quality of 20. Reads were excluded if they had multiple best hits or if they were tagged as not primary alignment, failure, or duplicate reads. The SFS was polarized using the ancestral state (see specific methods section on ancestral state generation). *realSFS* from *angsd* (*96*, *106*) was used to generate the 2D SFS between population pairs and generate $F_{ST}$ in sliding windows. Only windows with at least 100 SNPs were considered. $F_{ST}$ values were z-transformed and a cutoff of $z > 6$ was used for outlier windows. The polarised site frequency spectrum was used to generate neutrality tests in the same windows as $F_{ST}$ with the *thetaStat* tool in *angsd* (*112*). Fay & Wu's H (*113*) was extracted for $F_{ST}$-outlier and non-outlier windows and statistical significance was estimated with a Wilcoxon rank-sum test in *R*.

Gene ontology (GO) enrichment was assessed using GO terms from *A. thaliana* TAIR 10 (*80*) *BLAST* results. To identify GO terms enriched among candidate lists, the *topGO* package (*114*) in *R* (*115*) was used with Fisher's exact test, the *weight01* algorithm, and a $p < 0.05$ threshold to assess significance. Only GO terms describing biological processes were considered. In addition, $F_{ST}$ between individual SNPs was z-transformed and outliers were defined as those with $z > 100$. The annotation was used to examine if SNPs are located within gene regions. Functional annotation was obtained from *A. thaliana* TAIR 10 (*80*) *BLAST* results. Additionally, annotations were cross-referenced with 306 *A. thaliana* genes known to be involved in flowering time (*43*). 566 predicted *A. artemisiifolia* genes were matched to this dataset, representing 191 unique *A. thaliana* flowering time genes.

Metagenomic community classification

Reads that did not map against the *A. artemisiifolia* reference genome were used to analyse the leaf metagenome. Unmapped reads were extracted from the resulting BAM files using *samtools* v1.9 (*98*) and converted into *fastq* files with *PicardTools* v2.21.2 *SamToFastq* (http://broadinstitute.github.io/picard). Samples that were grown from seeds in the greenhouse were excluded from the analysis (see Data S4).

For the metagenomic classification, PCR duplicates were removed with *clumpify* from the *BBMap* package (sourceforge.net/projects/bbmap/) using the default parameters. The unmapped reads were then classified with *kraken2* (*116*) using the *NCBI_non-redundant_ntdb* database (downloaded August 2020) in paired-end mode for the paired data and in single-end mode for overlapping reads that were collapsed into a single read during adapter removal. As this study focuses on microbes, reads assigned to Metazoa or Viridiplantae were removed with the *extract_kraken_reads.py* script from *KrakenTools* (github.com/jenniferlu717/KrakenTools/). The two *kraken* reports for paired and collapsed reads for each sample were combined into one report using the *combine_kreports.py* script from *KrakenTools*. *Kraken-biom* v1.0.1 (github.com/smdabdoub/kraken-biom/) was used to combine the results from all samples for subsequent analysis in *R* v4.0.3.

Thereafter, several filters were applied to the dataset to exclude particular taxa from further analysis. Only taxonomic identifications at the species level were used. Species with a relative abundance below 0.05% were removed under the assumption that they are likely false positives. The package *decontam* (*117*) was used to identify taxa that are probable lab contaminants based on the extraction blanks using the *'prevalence'* method with a threshold of 0.4. Taxa identified as contamination, uncultured taxa and cloning vectors were removed. As we saw a large difference between historical herbarium and contemporary samples that we could not rule out as herbarium contamination, the analysis was restricted to prokaryotic plant pathogens identified by the *FAPROTAX* database (*118*) as these are less likely to be herbarium contamination. Abundance data was transformed into presence/absence, and significant differences were evaluated by using a two-sample t-test.

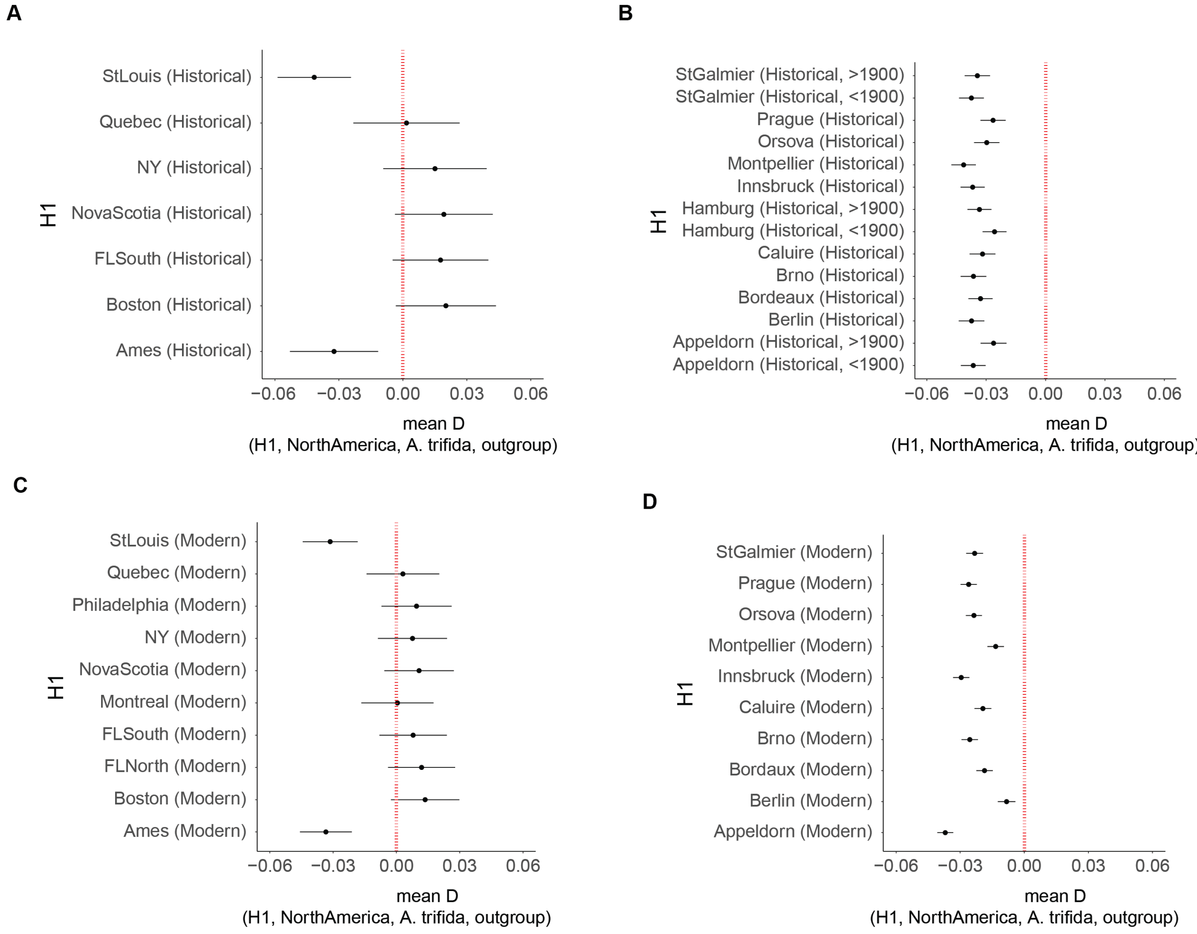


**Fig. S1. D statistics of the form (H1, North American population, *Ambrosia trifida*, *Ambrosia carduacea*). Plotted are the mean D values for all possible comparisons. Error bars indicate one standard deviation from the mean. (E) H1 and H2: Historical North American populations. (F) H1: Historical European populations, H2: Historical North American populations, excluding those that show signs of introgression with *A. trifida* (StLouis and Ames). (G) H1 and H2: Modern North American populations. (H) H1: Modern European populations, H2: Modern North American populations, excluding those that show signs of introgression with *A. trifida* (StLouis and Ames).**

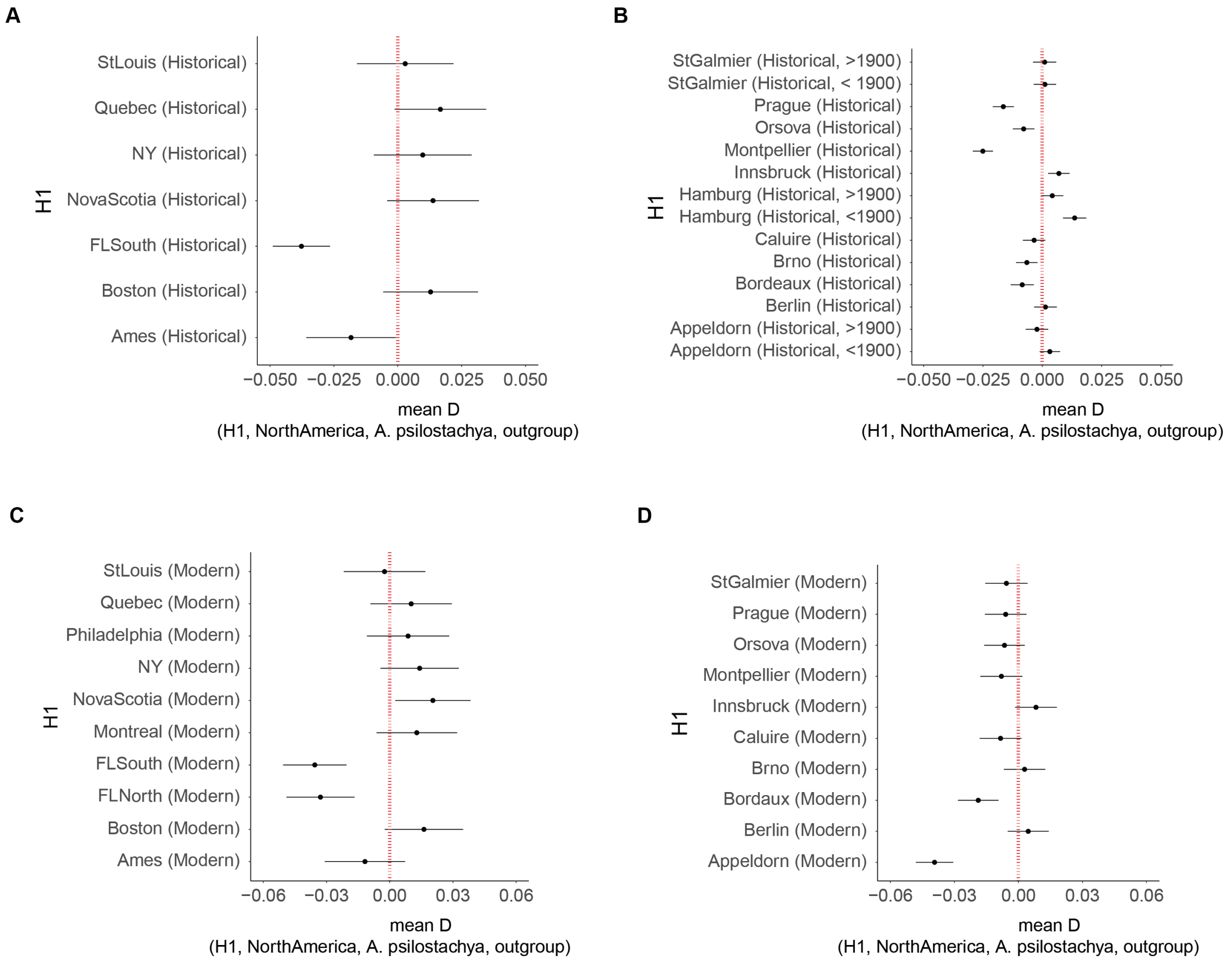


**Fig. S2. D statistics of the form (H1, North American population, *Ambrosia psilostachya*, *Ambrosia carduacea*). (A) H1 and H2: Historical North American populations. (B) H1: Historical European populations, H2: Historical North American populations, excluding those that show signs of introgression with *A. psilostachya* (FLSouth and Ames). (C) H1 and H2: Modern North American populations. (D) H1: Modern European populations, H2: Modern North American populations, excluding those that show signs of introgression with *A. psilostachya* (FLSouth and FLNorth).**

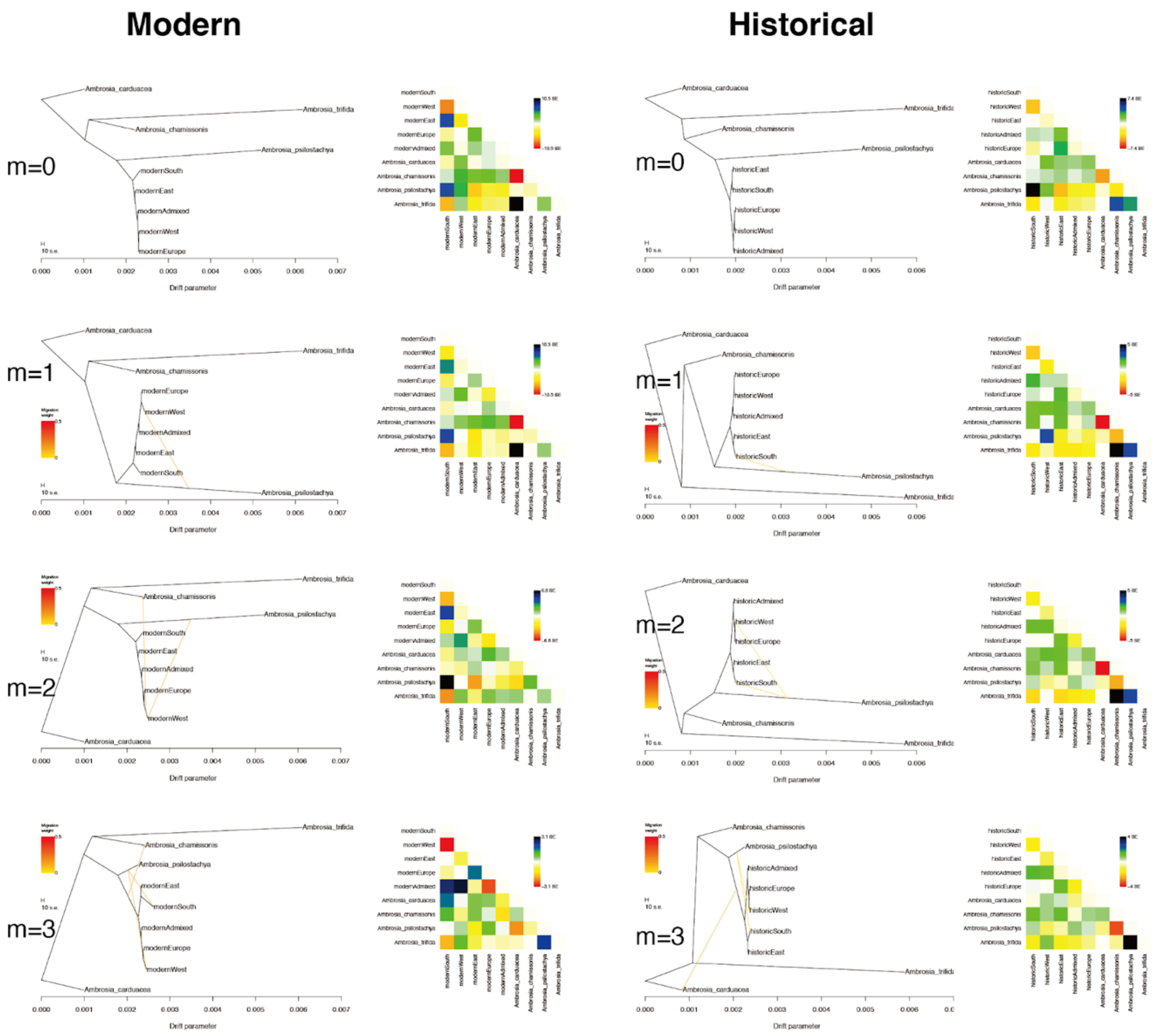


**Fig. S3. ML phylogeny reconstruction of modern and historical ragweed populations using Treemix. For modern and historical *Ambrosia artemisiifolia* populations, we randomly selected one allele per loci to get the population allele frequency. We fitted 0 to 3 (m=0-3) migration events indicated with arrows. Residual fit for each tree is shown right by the phylogeny. *Ambrosia carduacea* was used to root the tree.**

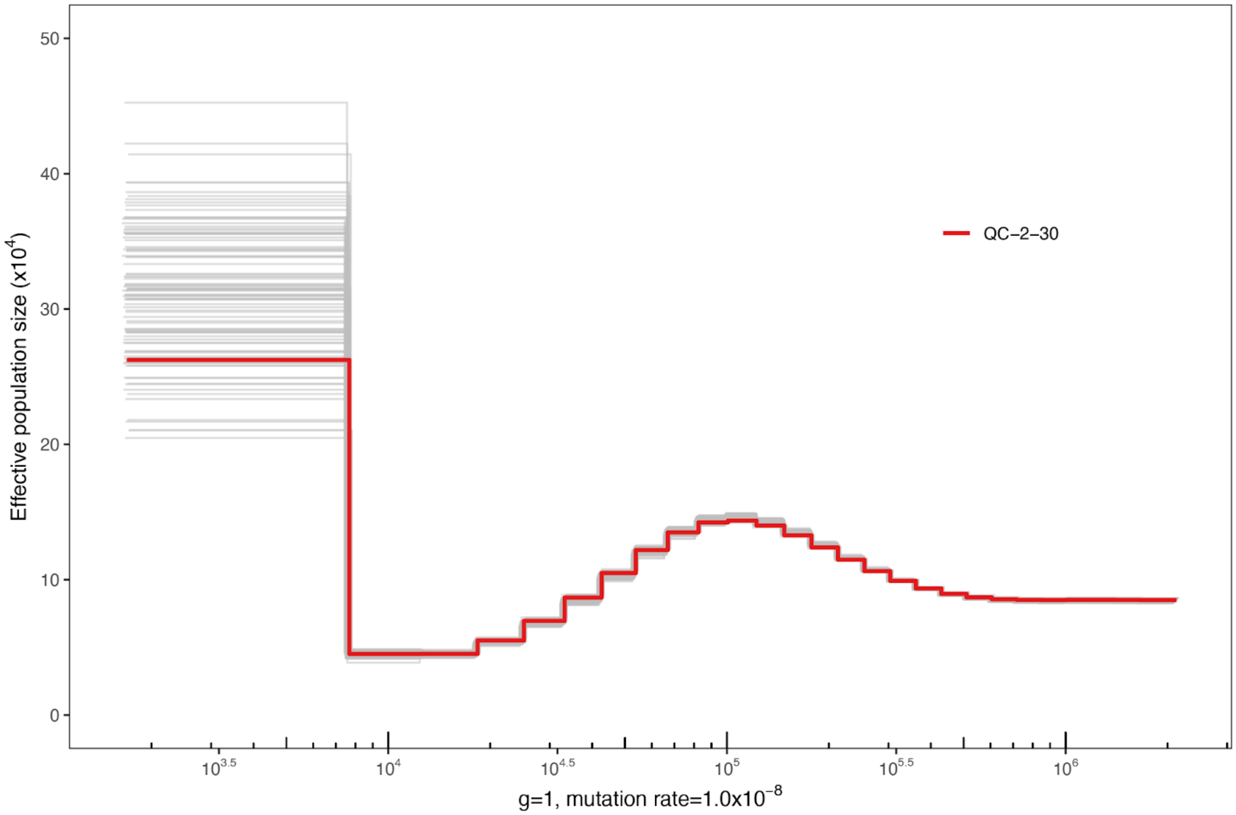


**Fig. S4. Demographic history of ragweed inferred by PSMC. We calibrated the effective population size assuming a mutation rate of 1.0 $10^{-8}$ per site per generation and the generation time of ragweed as 1 year. The grey lines represent 100 bootstraps. N. American individual QC-2-30 was used for demographic inference.**

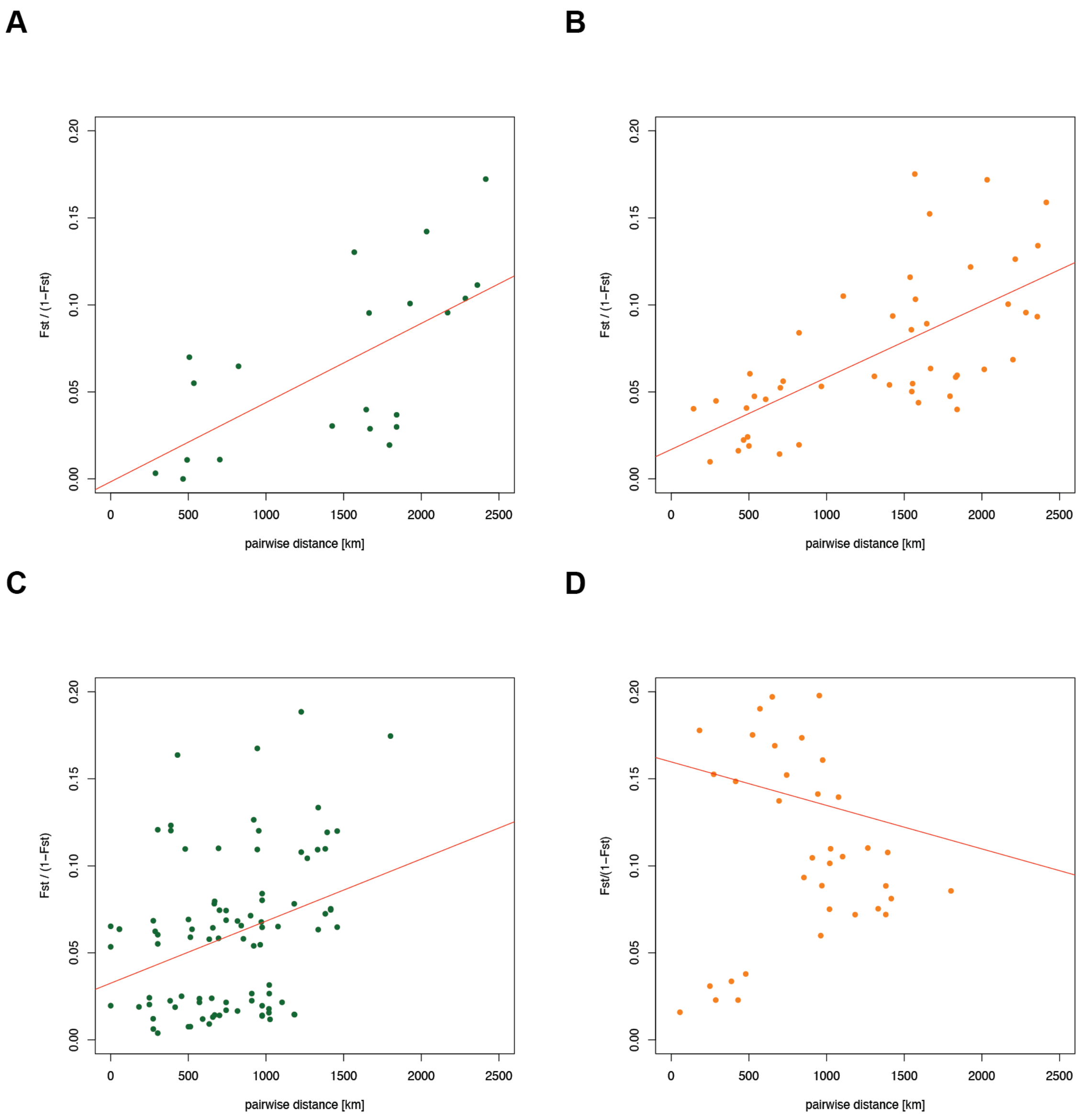


**Fig. S5. Isolation by distance in the native (top) and introduced (bottom) range. The red line represents the linear regression line. (A) historical North America (Mantel statistic r: 0.6462, significance: 0.004), (B) modern North America (Mantel statistic r: 0.6578,** significance: 0.001), (**C**) historical Europe (Mantel statistic r: 0.3244, significance: 0.051), (**D**) modern Europe (Mantel statistic r: -0.1234, significance: 0.696).

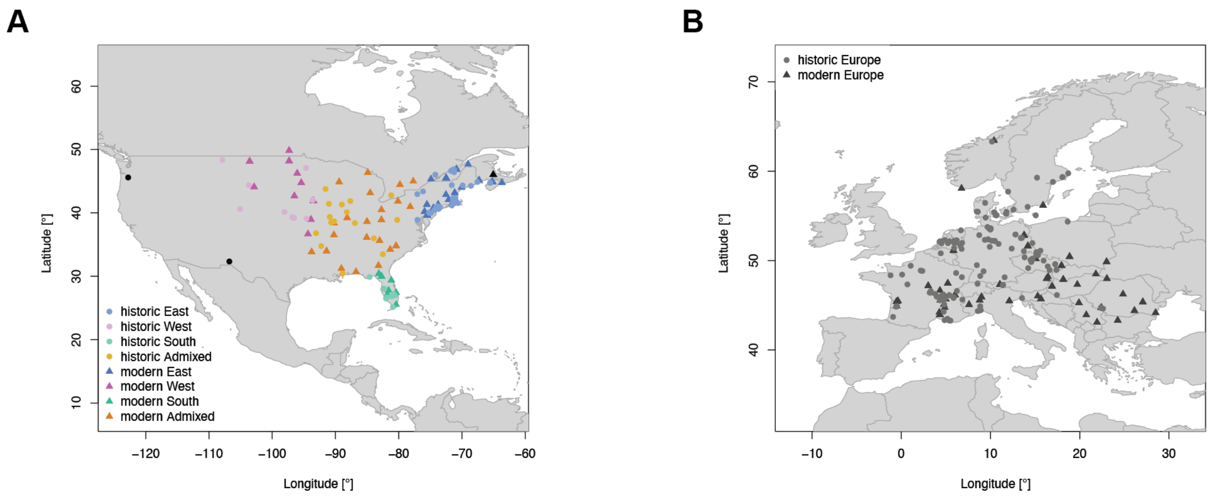


**Fig. S6. Sampling locations and population assignment. (A) North America. Light blue circles: historical East, dark blue triangles: modern East, light pink circles: historical West, dark pink triangles: modern West, light turquoise circles: historical South, dark turquoise triangles: modern South, light orange circles: historical Admixed, dark orange triangles: modern Admixed, black circles: historical samples that geographically do not group with their population and were thus excluded from population comparisons, black triangles: modern samples that geographically do not group with their population and were thus excluded from population comparisons. (B) Europe. grey circles: historical samples, black triangles: modern samples.**

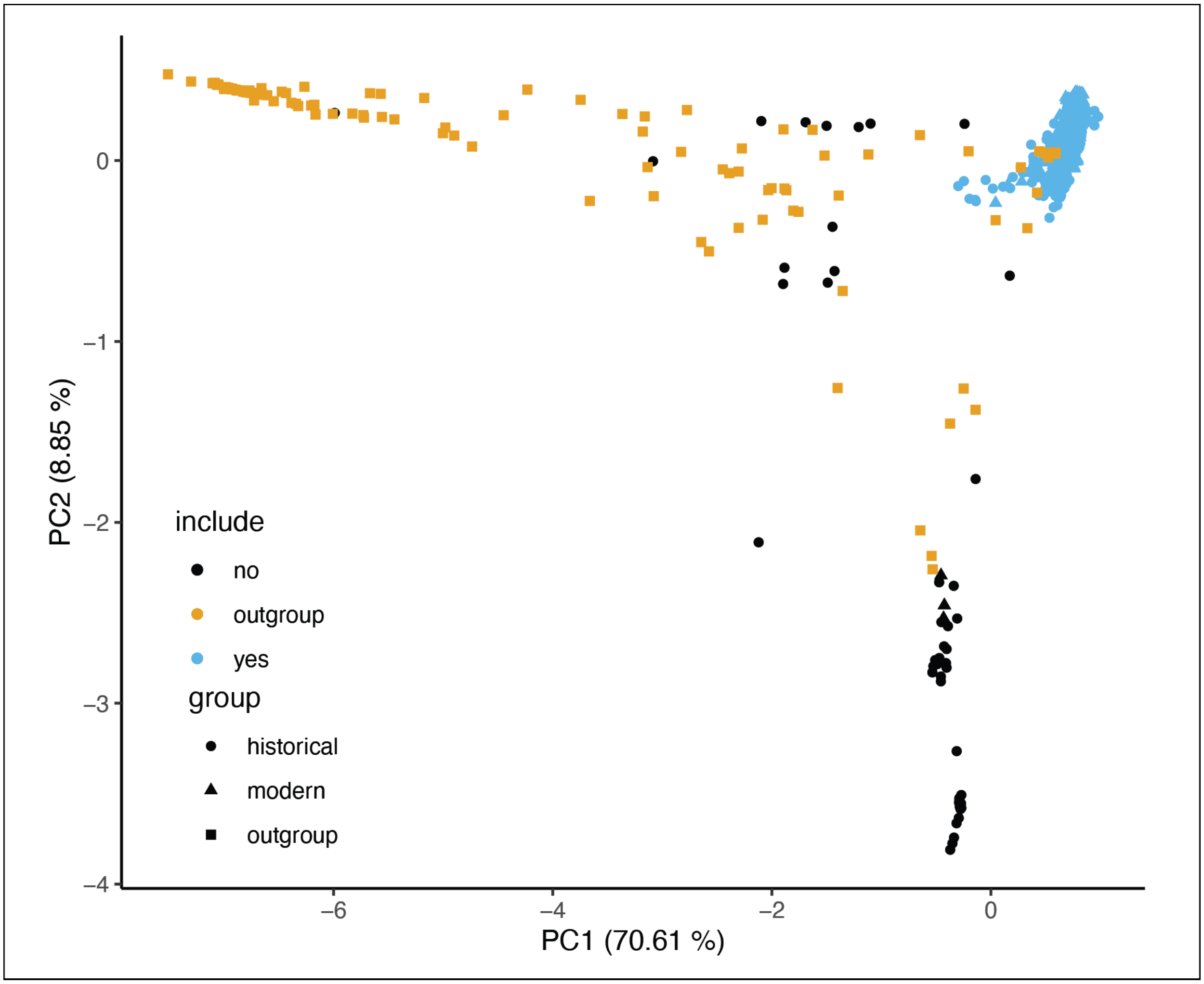


**Fig. S7. PCA including outgroup samples (other *Ambrosia* species). Orange squares: outgroup samples, black circles: historical samples that were excluded based on this PCA, blue circles: historical samples that were included in the final analysis, black triangles: modern samples that were excluded based on this PCA, blue triangles: modern samples that were included in the final analysis.**

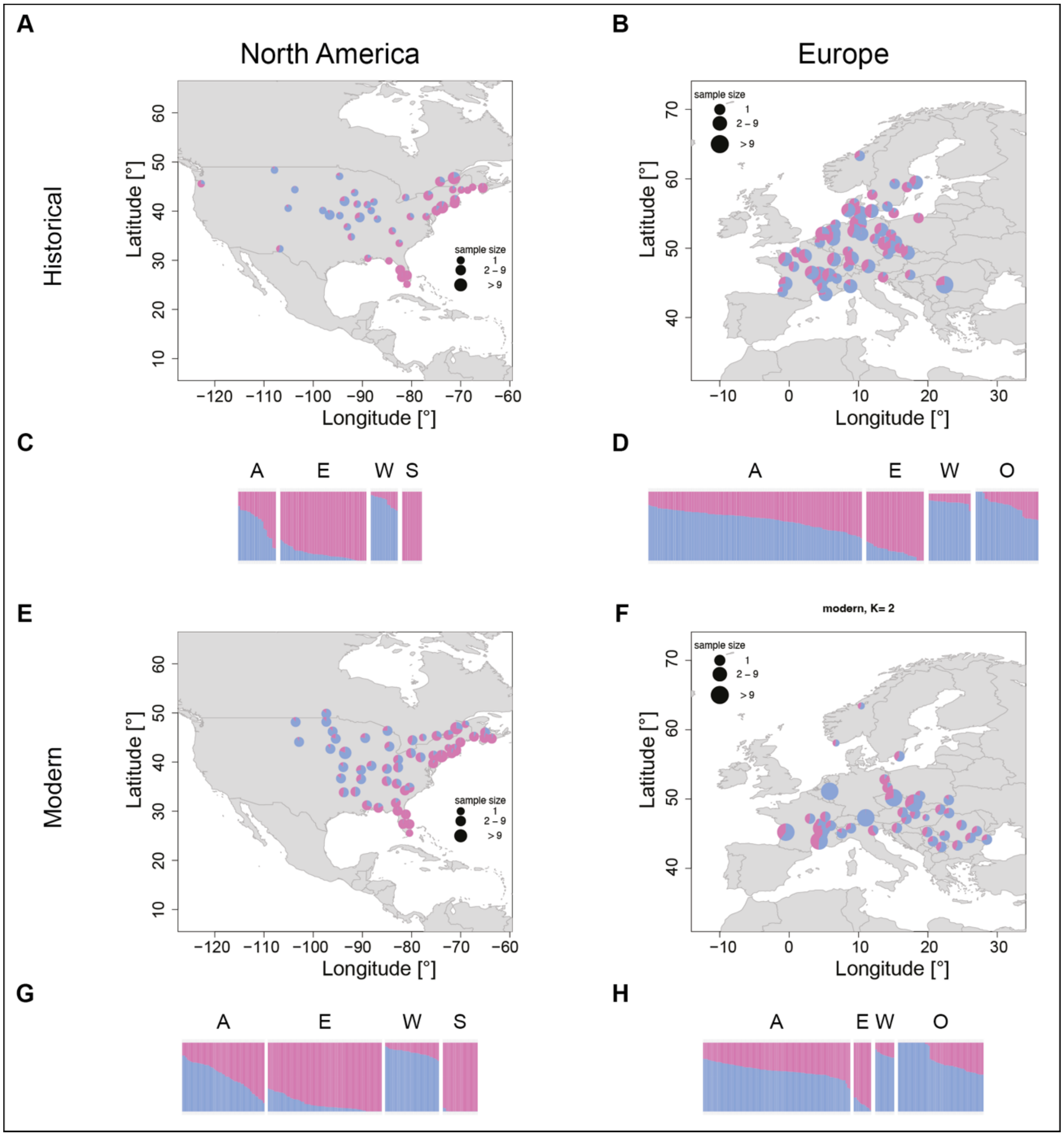


**Fig S8. Population structure obtained from NGSadmix for K=2. (A, B, E, F) Admixture maps. Samples within 100 km were grouped together and the average ancestry across those groups was plotted. If samples were grouped together, ancestry values were plotted at the centroid of the group. (C, D, G, H) Admixture barplots. Each bar represents one individual. Samples are grouped based on their assignment to a genetic cluster (based on K=9): A: Admixed, E: East, W: West, S: South, O: other. (A) Historical North America. (B) Historical Europe. (C) Historical North America. (D) Historical Europe. (E) Modern North America. (F) Modern Europe. (G) Modern North America. (H) Modern Europe. The NGSadmix run with the highest likelihood was used for plotting and the same color scheme was used across all panels.**

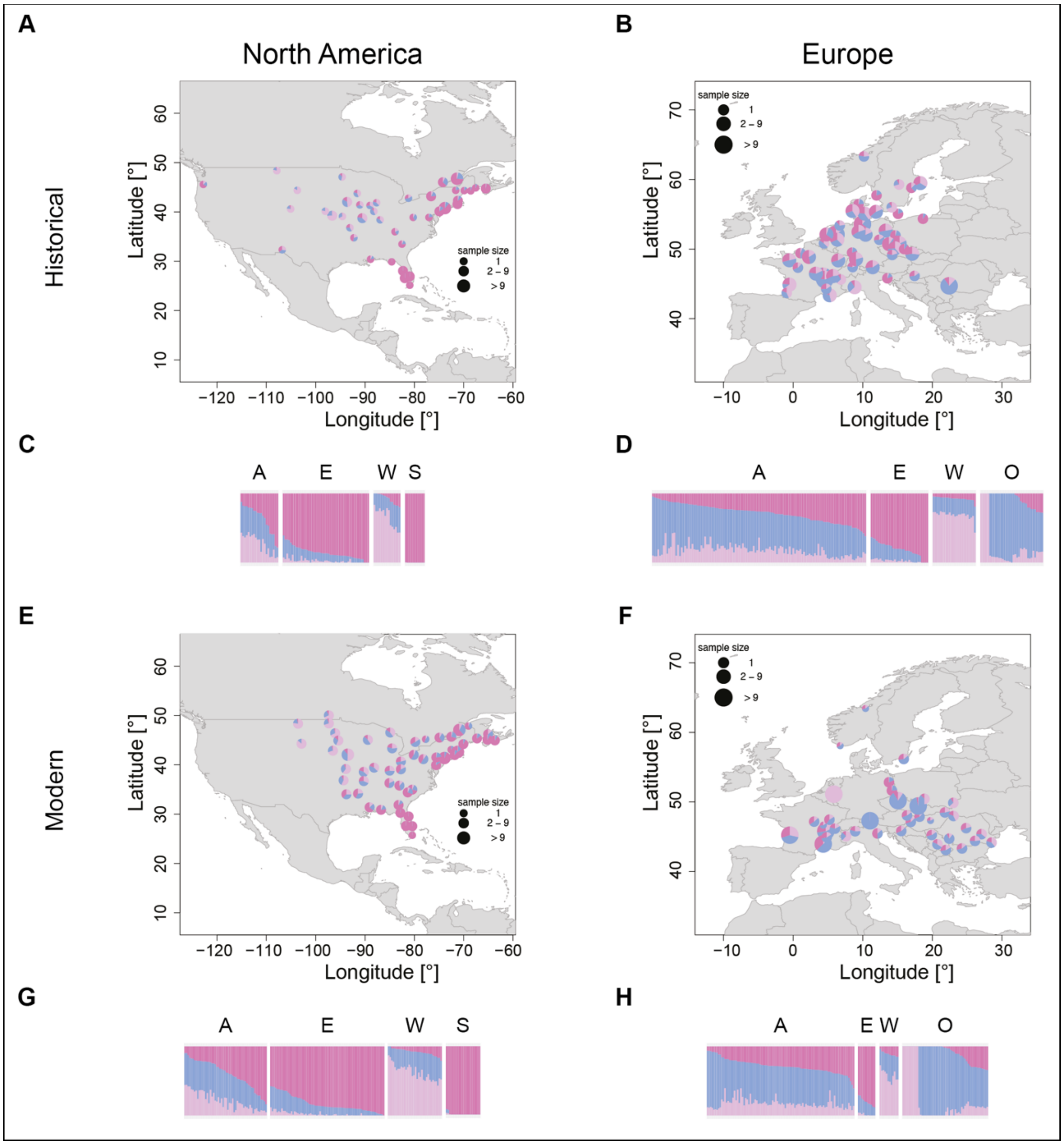


**Fig S9. Population structure obtained from NGSadmix for K=3. (A, B, E, F) Admixture maps. Samples within 100 km were grouped together and the average ancestry across those groups was plotted. If samples were grouped together, ancestry values were plotted at the centroid of the group. (C, D, G, H) Admixture barplots. Each bar represents one individual. Samples are grouped based on their assignment to a genetic cluster (based on K=9): A: Admixed, E: East, W: West, S: South, O: other. (A) Historical North America. (B) Historical Europe. (C) Historical North America. (D) Historical Europe. (E) Modern North America. (F) Modern Europe. (G) Modern North America. (H) Modern Europe. The NGSadmix run with the highest likelihood was used for plotting and the same color scheme was used across all panels.**

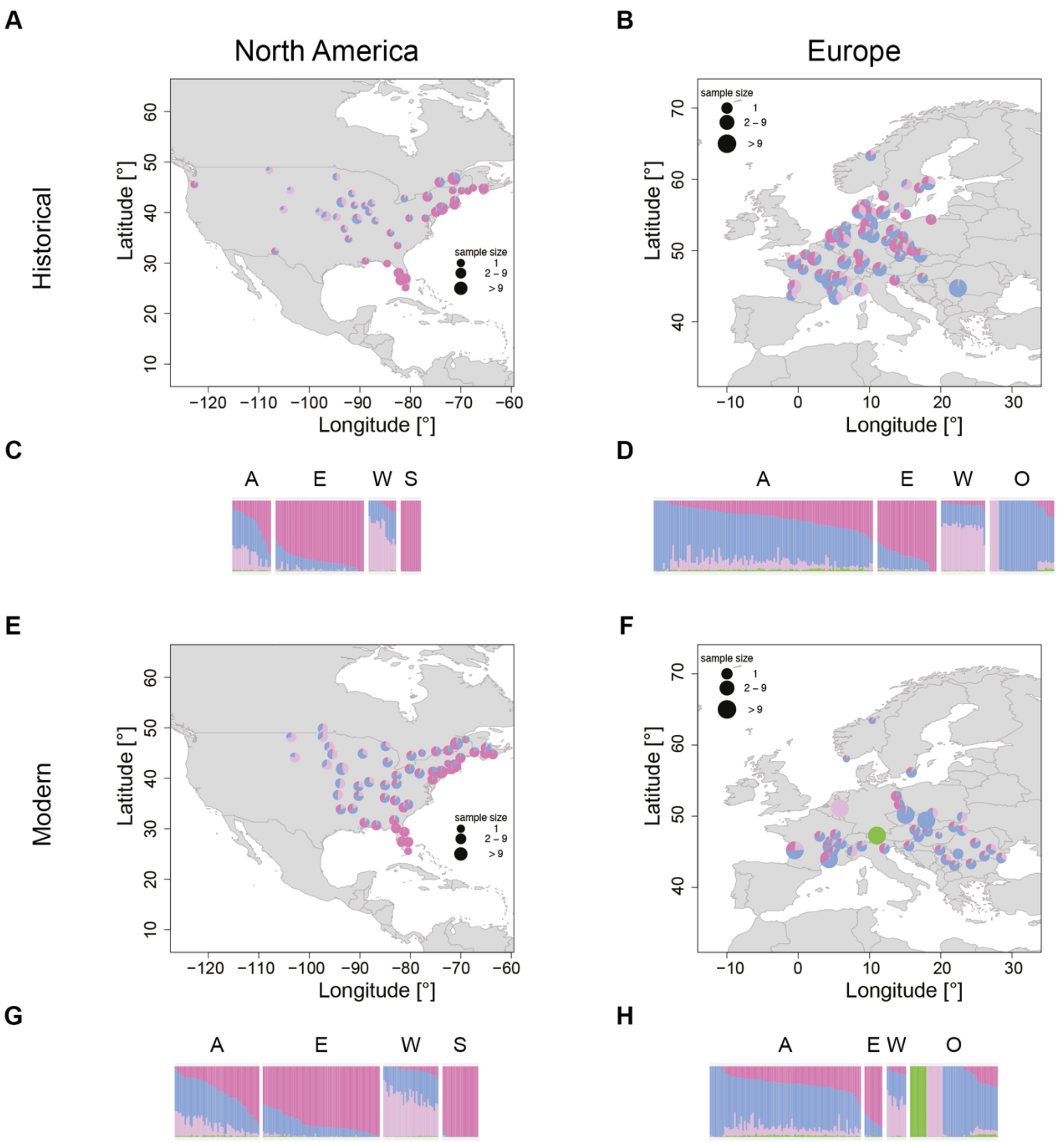


**Fig S10. Population structure obtained from NGSadmix for K=4. (A, B, E, F) Admixture maps. Samples within 100 km were grouped together and the average ancestry across those groups was plotted. If samples were grouped together, ancestry values were plotted at the centroid of the group. (C, D, G, H) Admixture barplots. Each bar represents one individual. Samples are grouped based on their assignment to a genetic cluster (based on K=9): A: Admixed, E: East, W: West, S: South, O: other. (A) Historical North America. (B) Historical Europe. (C) Historical North America. (D) Historical Europe. (E) Modern North America. (F) Modern Europe. (G) Modern North America. (H) Modern Europe. The NGSadmix run with the highest likelihood was used for plotting and the same color scheme was used across all panels.**

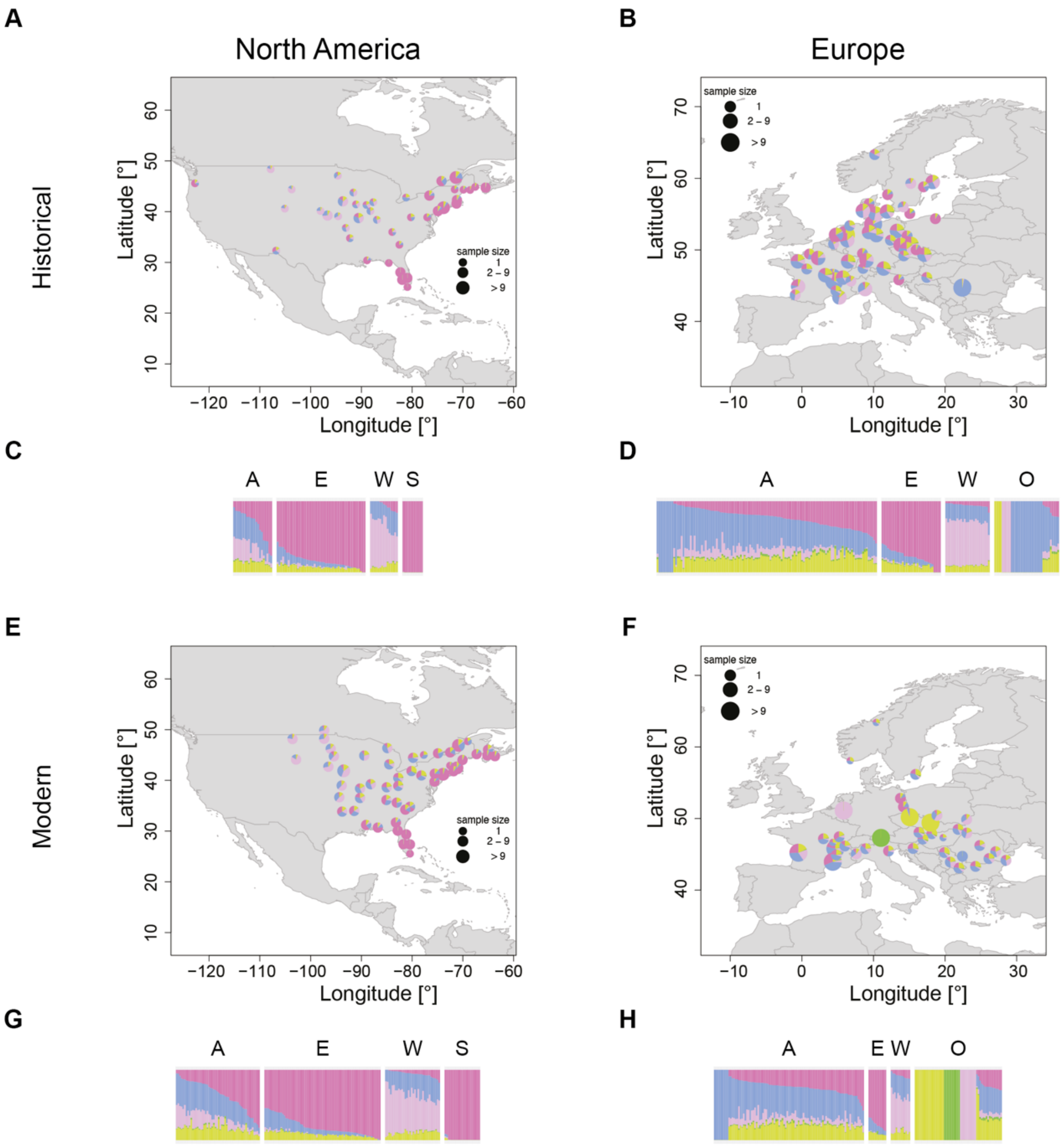


**Fig S11. Population structure obtained from NGSadmix for K=5. (A, B, E, F) Admixture maps. Samples within 100 km were grouped together and the average ancestry across those groups was plotted. If samples were grouped together, ancestry values were plotted at the centroid of the group. (C, D, G, H) Admixture barplots. Each bar represents one individual. Samples are grouped based on their assignment to a genetic cluster (based on K=9): A: Admixed, E: East, W: West, S: South, O: other. (A) Historical North America. (B) Historical Europe. (C) Historical North America. (D) Historical Europe. (E) Modern North America. (F) Modern Europe. (G) Modern North America. (H) Modern Europe. The NGSadmix run with the highest likelihood was used for plotting and the same color scheme was used across all panels.**

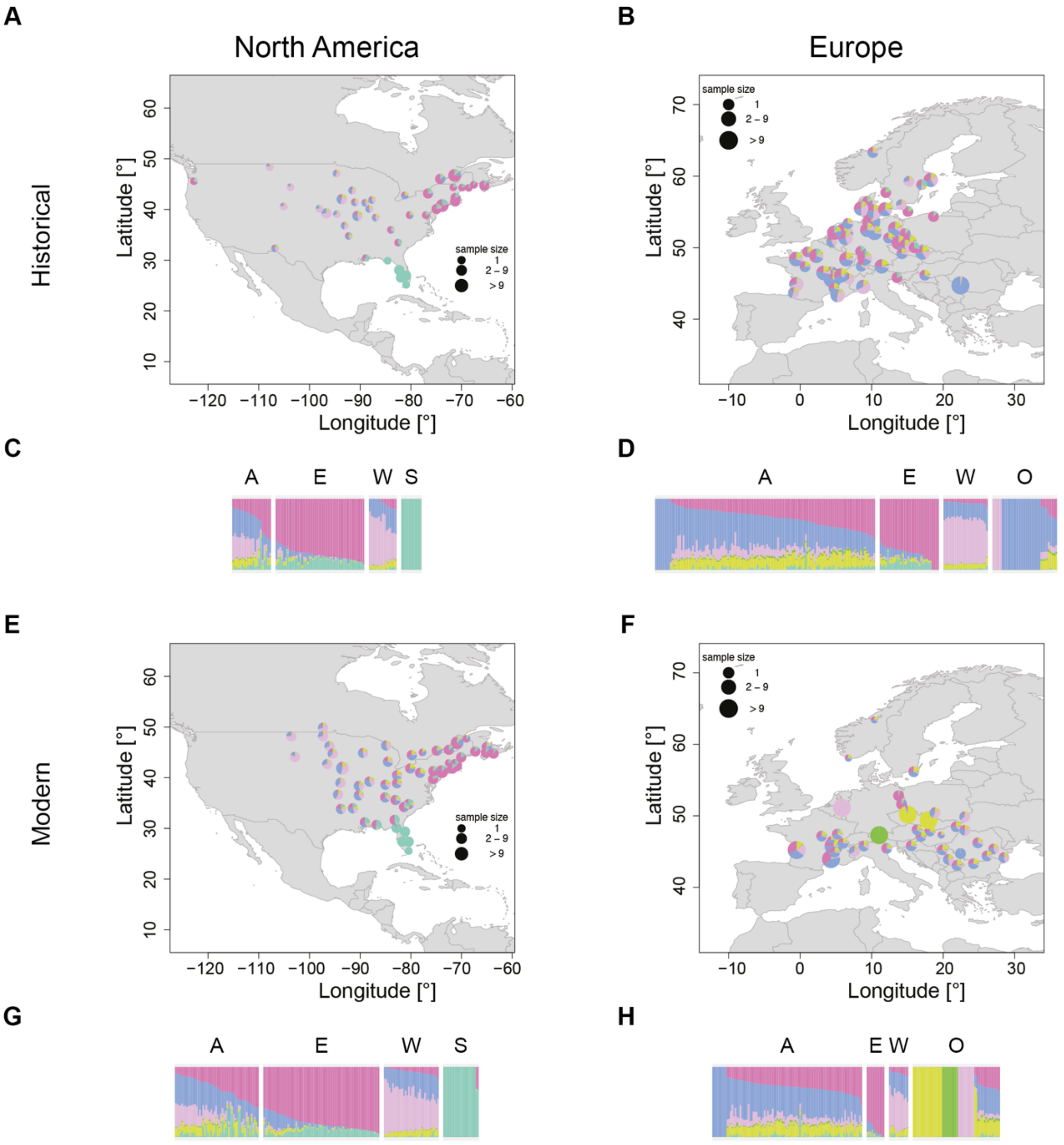


**Fig S12. Population structure obtained from NGSadmix for K=6. (A, B, E, F) Admixture maps. Samples within 100 km were grouped together and the average ancestry across those groups was plotted. If samples were grouped together, ancestry values were plotted at the centroid of the group. (C, D, G, H) Admixture barplots. Each bar represents one individual. Samples are grouped based on their assignment to a genetic cluster (based on K=9): A: Admixed, E: East, W: West, S: South, O: other. (A) Historical North America. (B) Historical Europe. (C) Historical North America. (D) Historical Europe. (E) Modern North America. (F) Modern Europe. (G) Modern North America. (H) Modern Europe. The NGSadmix run with the highest likelihood was used for plotting and the same color scheme was used across all panels.**

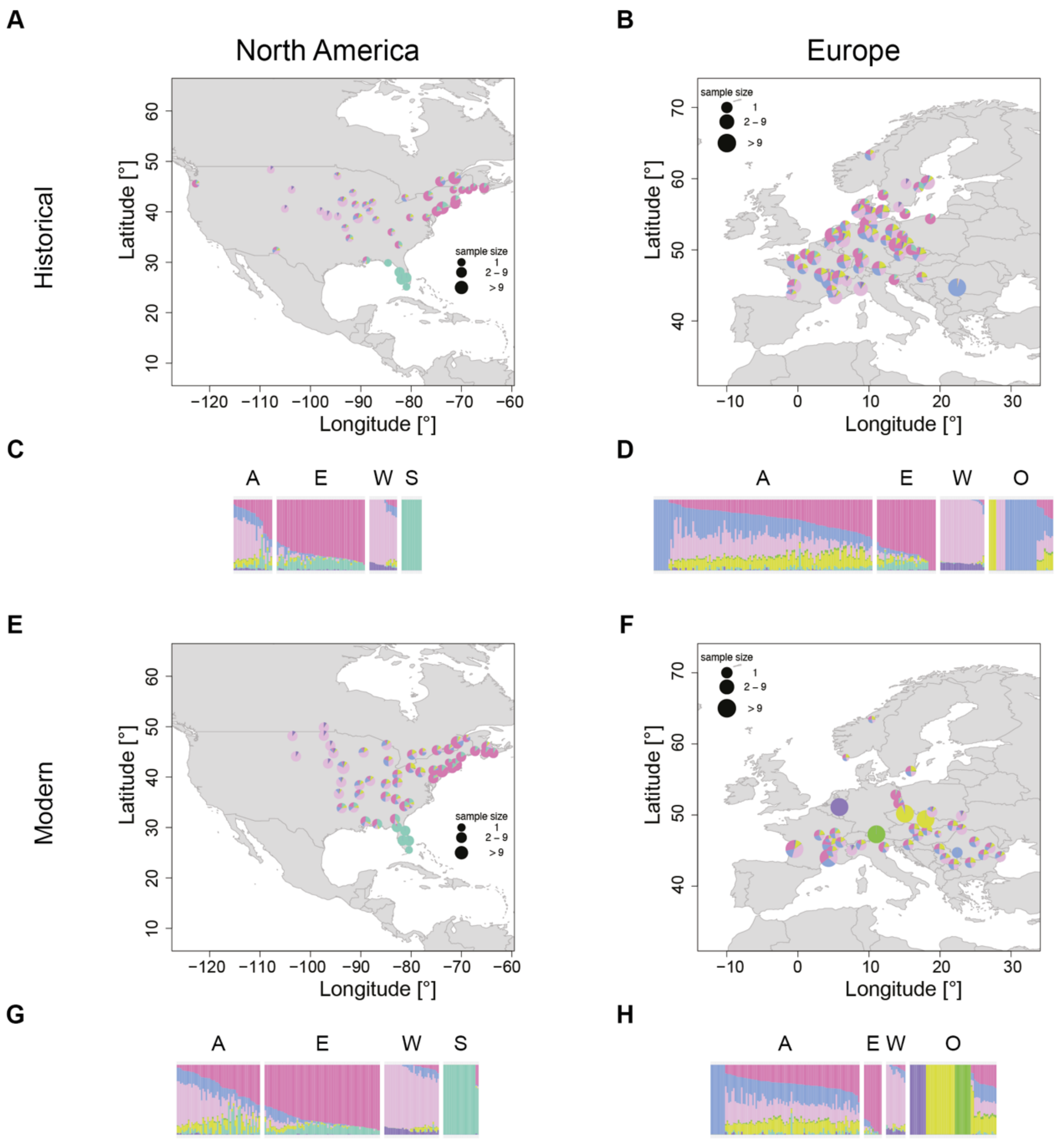


**Fig S13. Population structure obtained from NGSadmix for K=7. (A, B, E, F) Admixture maps. Samples within 100 km were grouped together and the average ancestry across those groups was plotted. If samples were grouped together, ancestry values were plotted at the centroid of the group. (C, D, G, H) Admixture barplots. Each bar represents one individual. Samples are grouped based on their assignment to a genetic cluster (based on K=9): A: Admixed, E: East, W: West, S: South, O: other. (A) Historical North America. (B) Historical Europe. (C) Historical North America. (D) Historical Europe. (E) Modern North America. (F) Modern Europe. (G) Modern North America. (H) Modern Europe. The NGSadmix run with the highest likelihood was used for plotting and the same color scheme was used across all panels.**

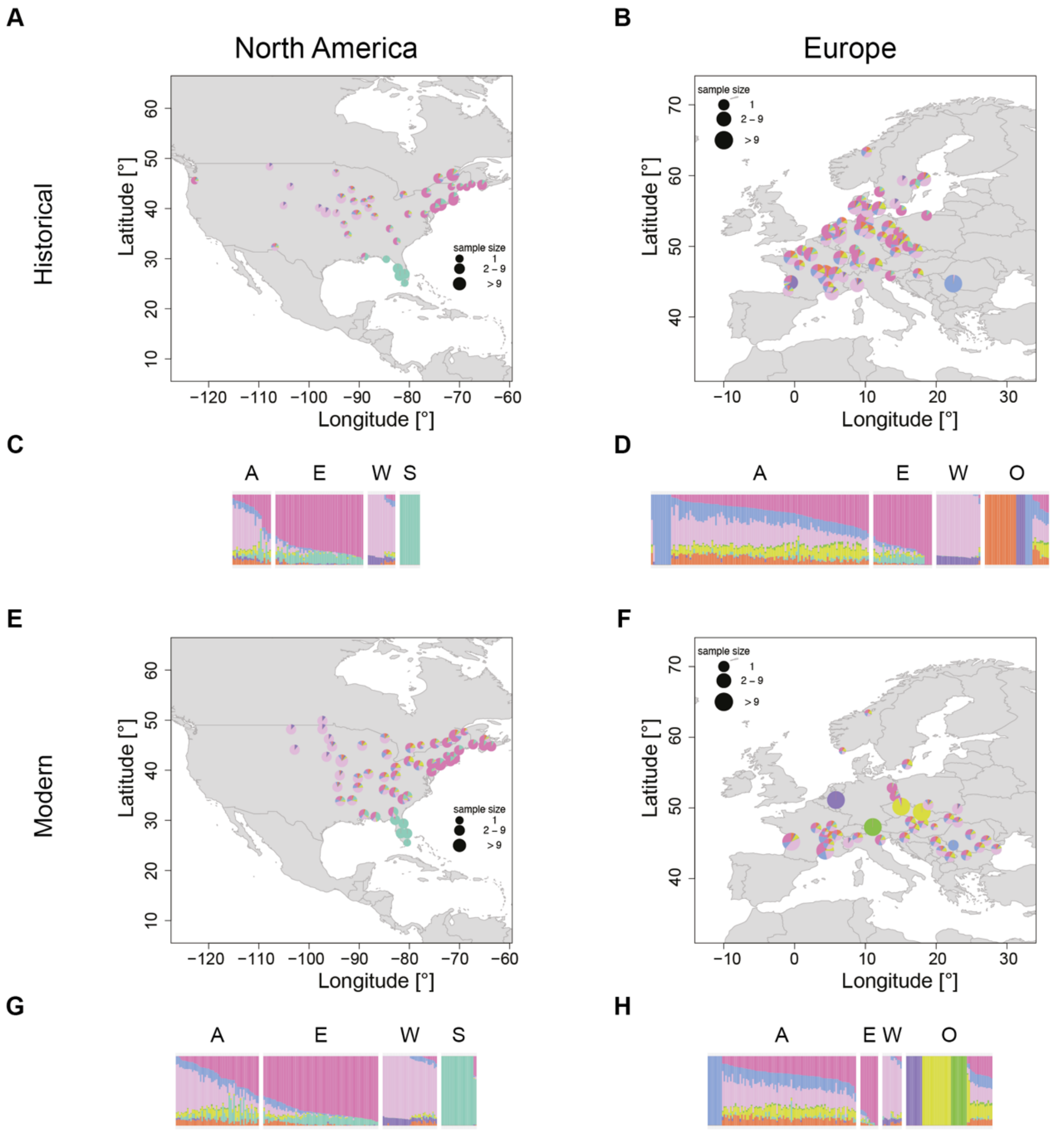


**Fig S14. Population structure obtained from NGSadmix for K=8. (A, B, E, F) Admixture maps. Samples within 100 km were grouped together and the average ancestry across those groups was plotted. If samples were grouped together, ancestry values were plotted at the centroid of the group. (C, D, G, H) Admixture barplots. Each bar represents one individual. Samples are grouped based on their assignment to a genetic cluster (based on K=9): A: Admixed, E: East, W: West, S: South, O: other. (A) Historical North America. (B) Historical Europe. (C) Historical North America. (D) Historical Europe. (E) Modern North America. (F) Modern Europe. (G) Modern North America. (H) Modern Europe. The NGSadmix run with the highest likelihood was used for plotting and the same color scheme was used across all panels.**

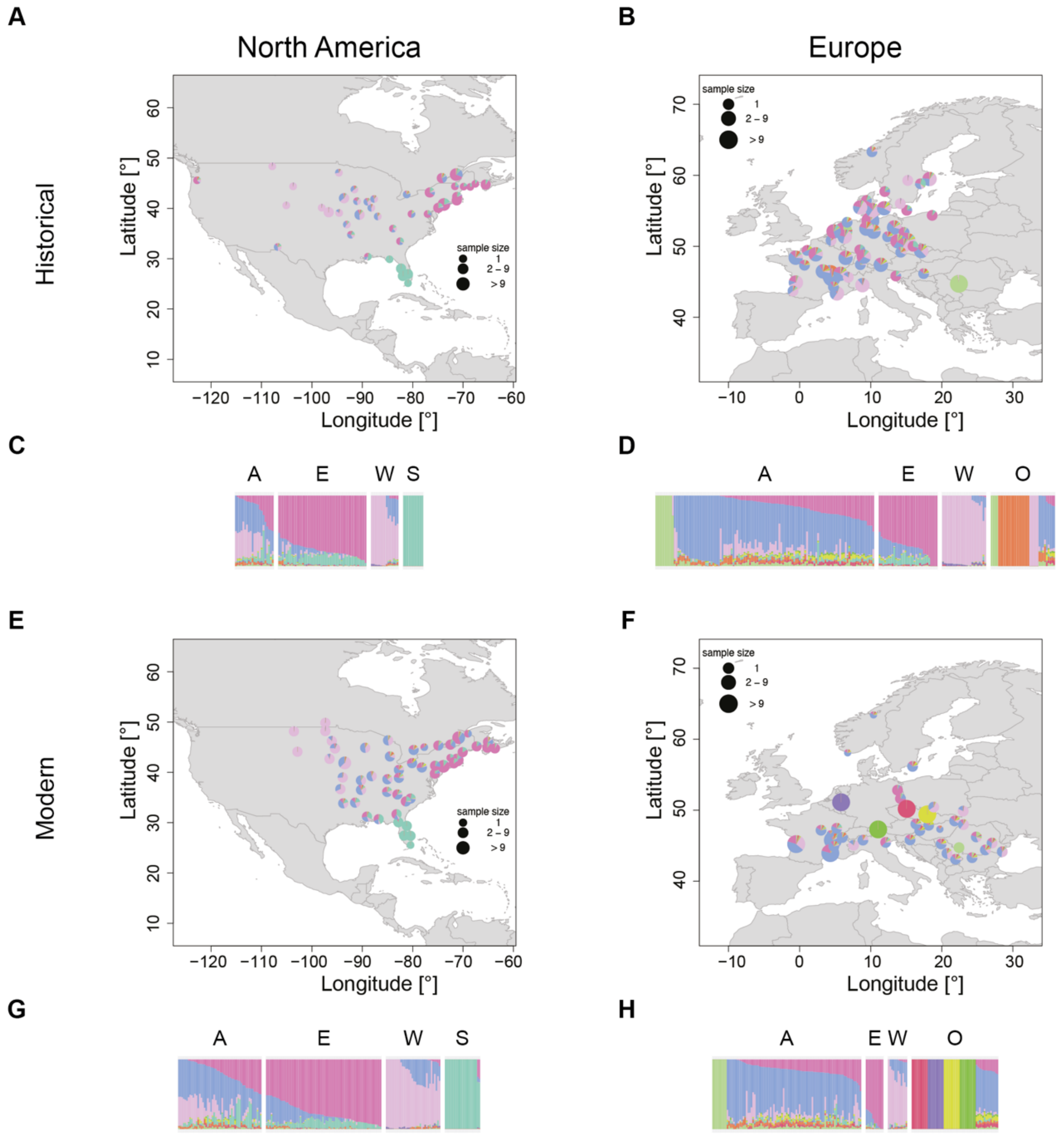


**Fig S15. Population structure obtained from NGSadmix for K=10. (A, B, E, F) Admixture maps. Samples within 100 km were grouped together and the average ancestry across those groups was plotted. If samples were grouped together, ancestry values were plotted at the centroid of the group. (C, D, G, H) Admixture barplots. Each bar represents one individual. Samples are grouped based on their assignment to a genetic cluster (based on K=9): A: Admixed, E: East, W: West, S: South, O: other. (A) Historical North America. (B) Historical Europe. (C) Historical North America. (D) Historical Europe. (E) Modern North America. (F) Modern Europe. (G) Modern North America. (H) Modern Europe. The NGSadmix run with the highest likelihood was used for plotting and the same color scheme was used across all panels.**

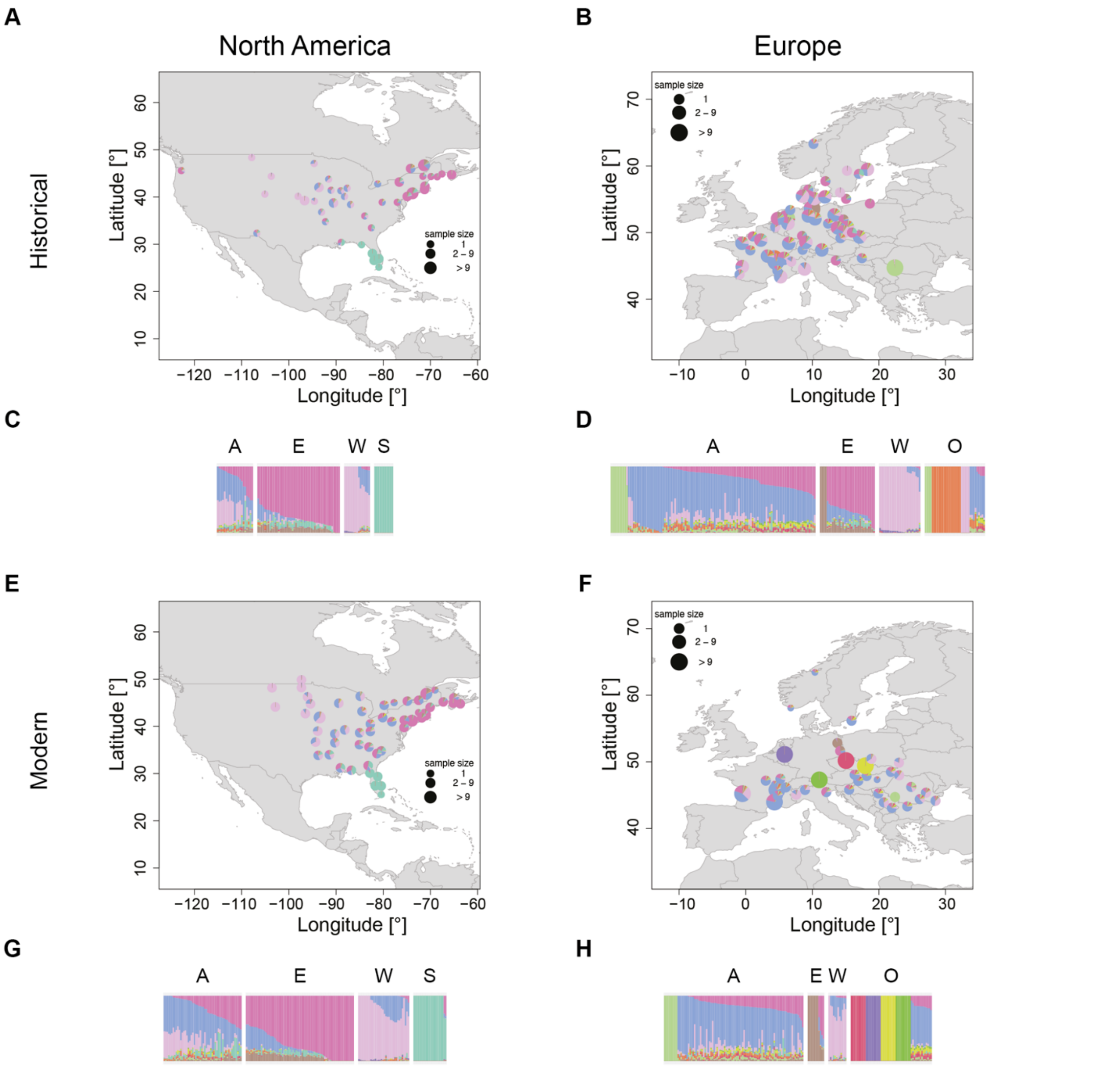


**Fig S16. Population structure obtained from NGSadmix for K=11. (A, B, E, F) Admixture maps. Samples within 100 km were grouped together and the average ancestry across those groups was plotted. If samples were grouped together, ancestry values were plotted at the centroid of the group. (C, D, G, H) Admixture barplots. Each bar represents one individual. Samples are grouped based on their assignment to a genetic cluster (based on K=9): A: Admixed, E: East, W: West, S: South, O: other. (A) Historical North America. (B) Historical Europe. (C) Historical North America. (D) Historical Europe. (E) Modern North America. (F) Modern Europe. (G) Modern North America. (H) Modern Europe. The NGSadmix run with the highest likelihood was used for plotting and the same color scheme was used across all panels.**

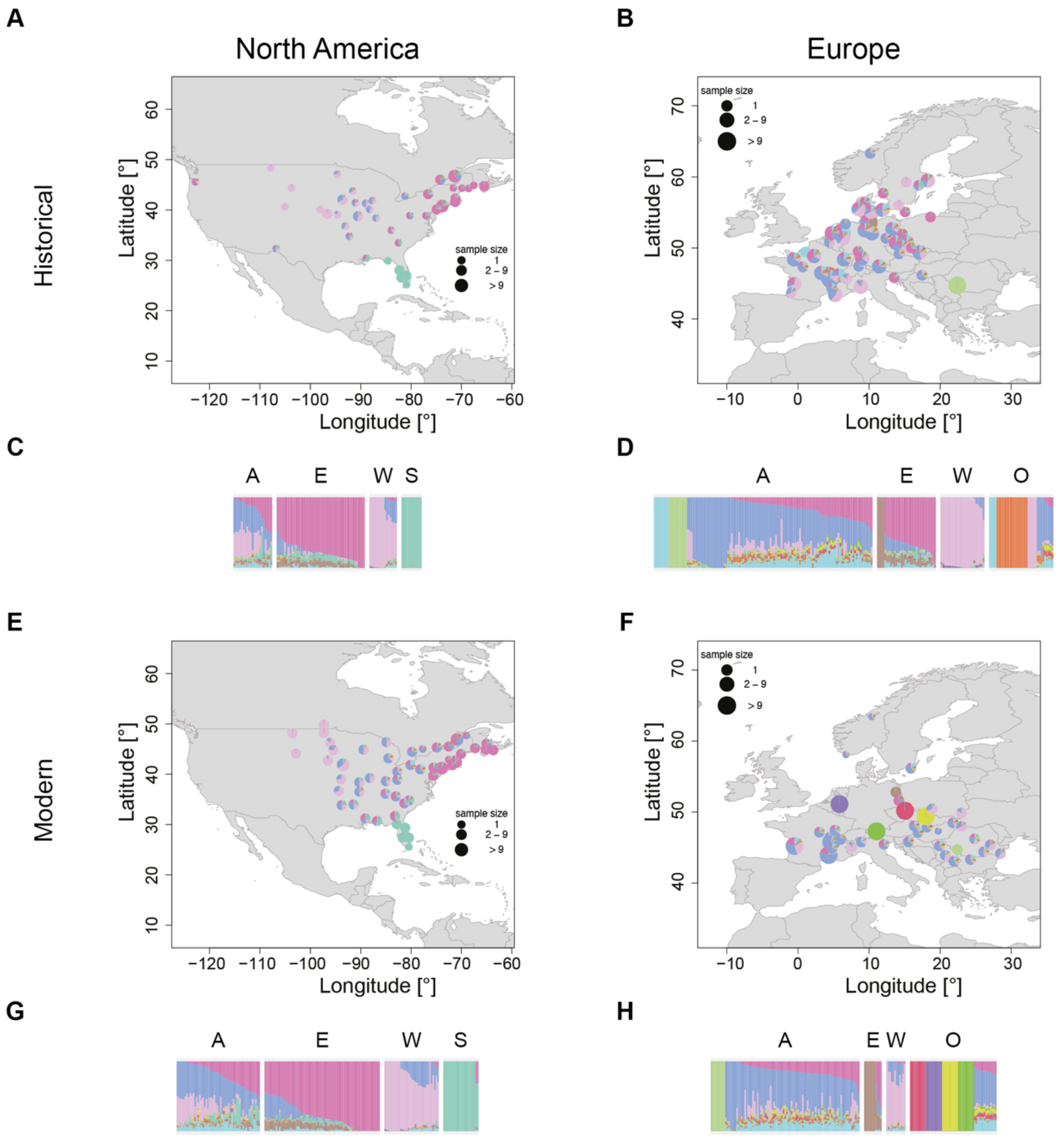


**Fig S17. Population structure obtained from NGSadmix for K=12. (A, B, E, F) Admixture maps. Samples within 100 km were grouped together and the average ancestry across those groups was plotted. If samples were grouped together, ancestry values were plotted at the centroid of the group. (C, D, G, H) Admixture barplots. Each bar represents one individual. Samples are grouped based on their assignment to a genetic cluster (based on K=9): A: Admixed, E: East, W: West, S: South, O: other. (A) Historical North America. (B) Historical Europe. (C) Historical North America. (D) Historical Europe. (E) Modern North America. (F) Modern Europe. (G) Modern North America. (H) Modern Europe. The NGSadmix run with the highest likelihood was used for plotting and the same color scheme was used across all panels.**

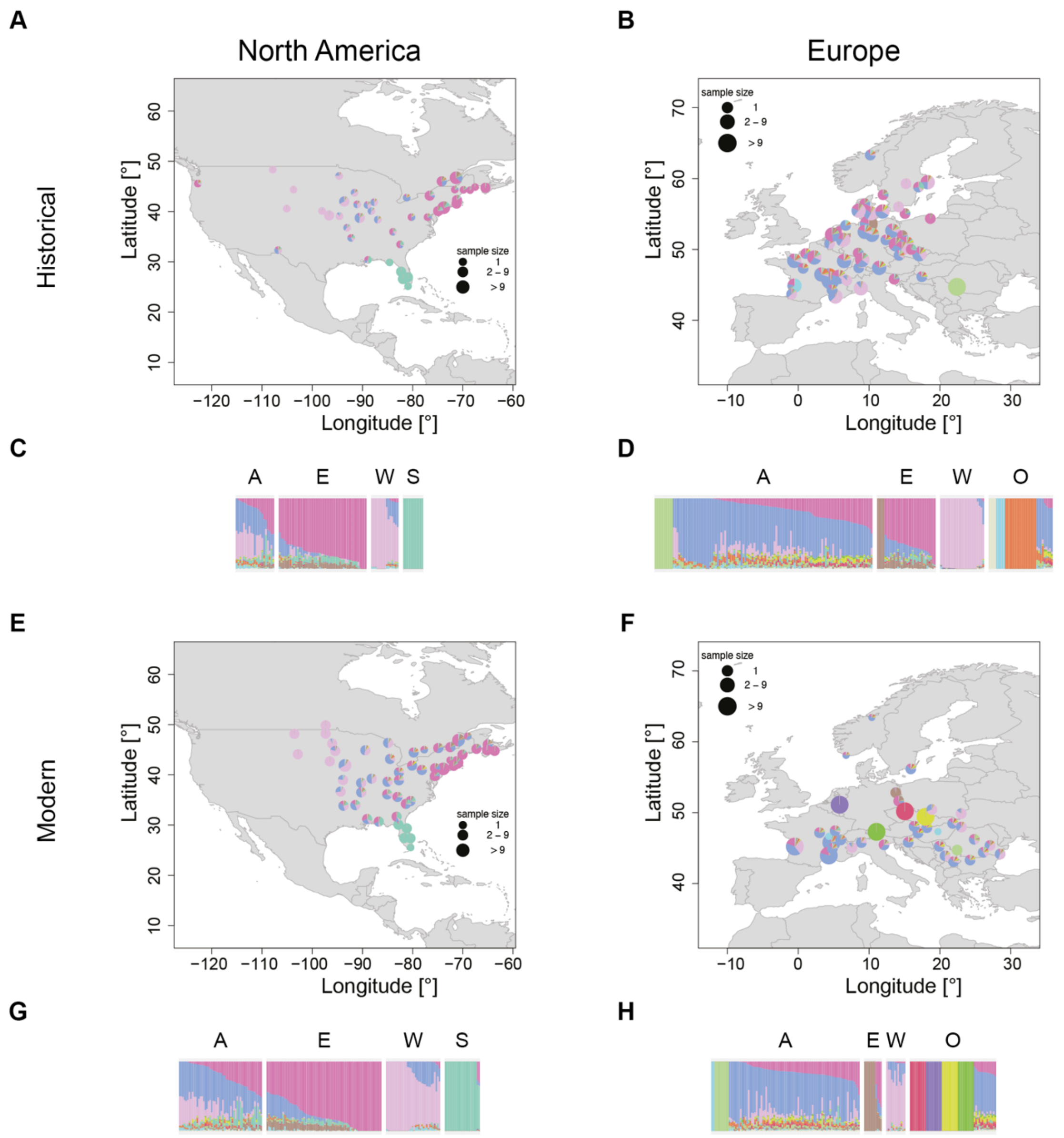


**Fig S18. Population structure obtained from NGSadmix for K=13. (A, B, E, F) Admixture maps. Samples within 100 km were grouped together and the average ancestry across those groups was plotted. If samples were grouped together, ancestry values were plotted at the centroid of the group. (C, D, G, H) Admixture barplots. Each bar represents one individual. Samples are grouped based on their assignment to a genetic cluster (based on K=9): A: Admixed, E: East, W: West, S: South, O: other. (A) Historical North America. (B) Historical Europe. (C) Historical North America. (D) Historical Europe. (E) Modern North America. (F) Modern Europe. (G) Modern North America. (H) Modern Europe. The NGSadmix run with the highest likelihood was used for plotting and the same color scheme was used across all panels.**

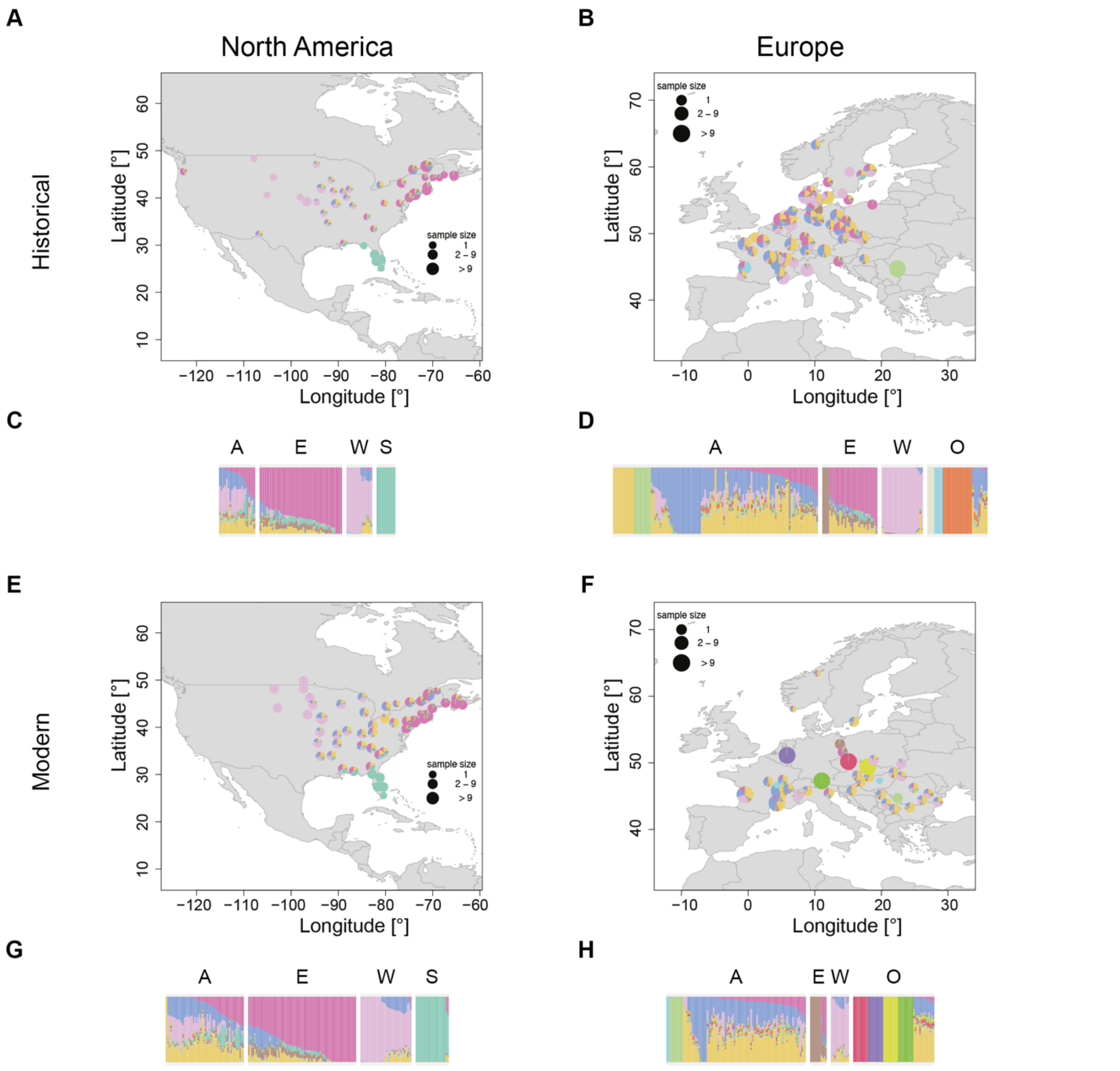


**Fig S19. Population structure obtained from NGSadmix for K=14. (A, B, E, F) Admixture maps. Samples within 100 km were grouped together and the average ancestry across those groups was plotted. If samples were grouped together, ancestry values were plotted at the centroid of the group. (C, D, G, H) Admixture barplots. Each bar represents one individual. Samples are grouped based on their assignment to a genetic cluster (based on K=9): A: Admixed, E: East, W: West, S: South, O: other. (A) Historical North America. (B) Historical Europe. (C) Historical North America. (D) Historical Europe. (E) Modern North America. (F) Modern Europe. (G) Modern North America. (H) Modern Europe. The NGSadmix run with the highest likelihood was used for plotting and the same color scheme was used across all panels.**

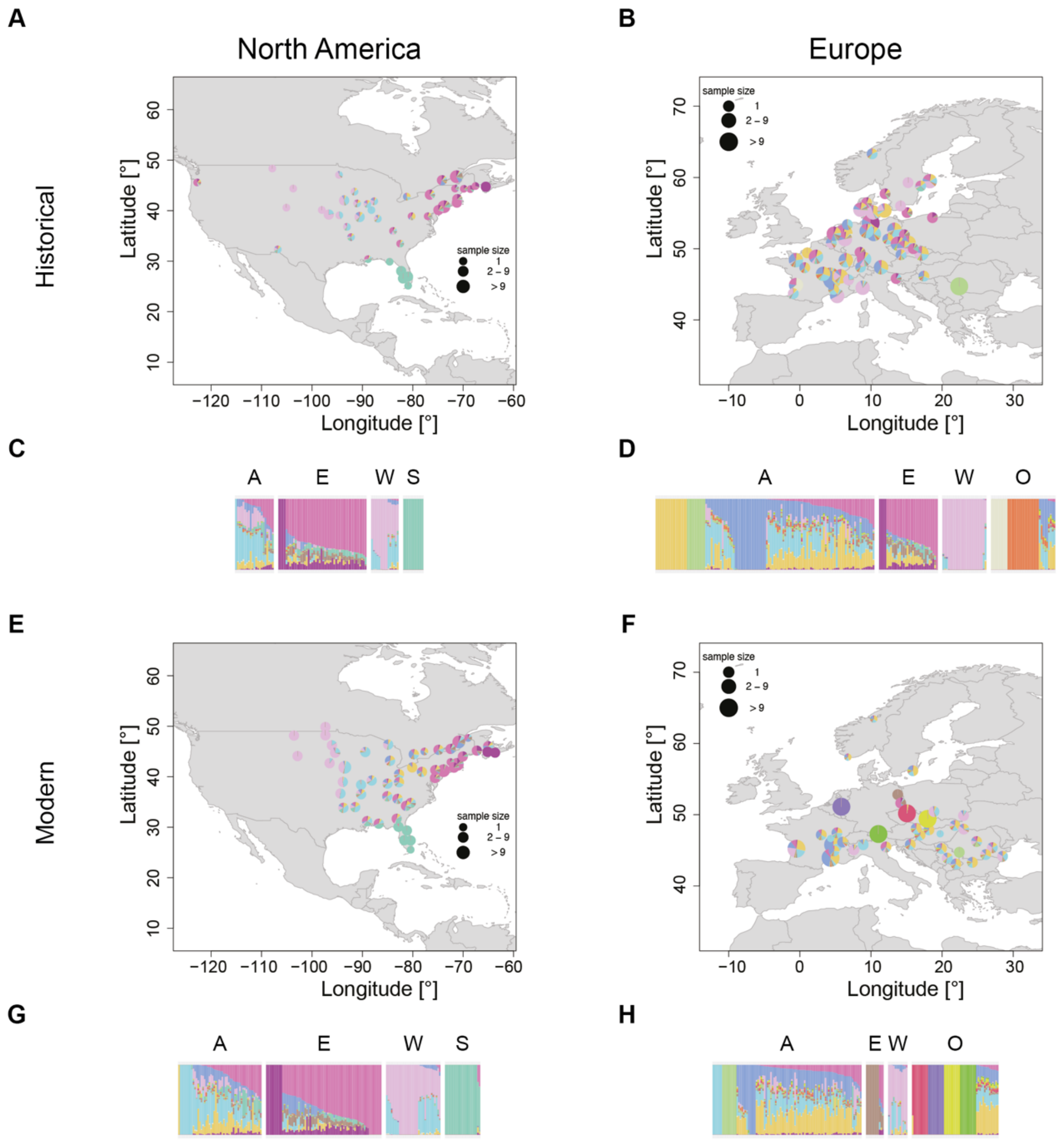


**Fig S20. Population structure obtained from NGSadmix for K=15. (A, B, E, F) Admixture maps. Samples within 100 km were grouped together and the average ancestry across those groups was plotted. If samples were grouped together, ancestry values were plotted at the centroid of the group. (C, D, G, H) Admixture barplots. Each bar represents one individual. Samples are grouped based on their assignment to a genetic cluster (based on K=9): A: Admixed, E: East, W: West, S: South, O: other. (A) Historical North America. (B) Historical Europe. (C) Historical North America. (D) Historical Europe. (E) Modern North America. (F) Modern Europe. (G) Modern North America. (H) Modern Europe. The NGSadmix run with the highest likelihood was used for plotting and the same color scheme was used across all panels.**

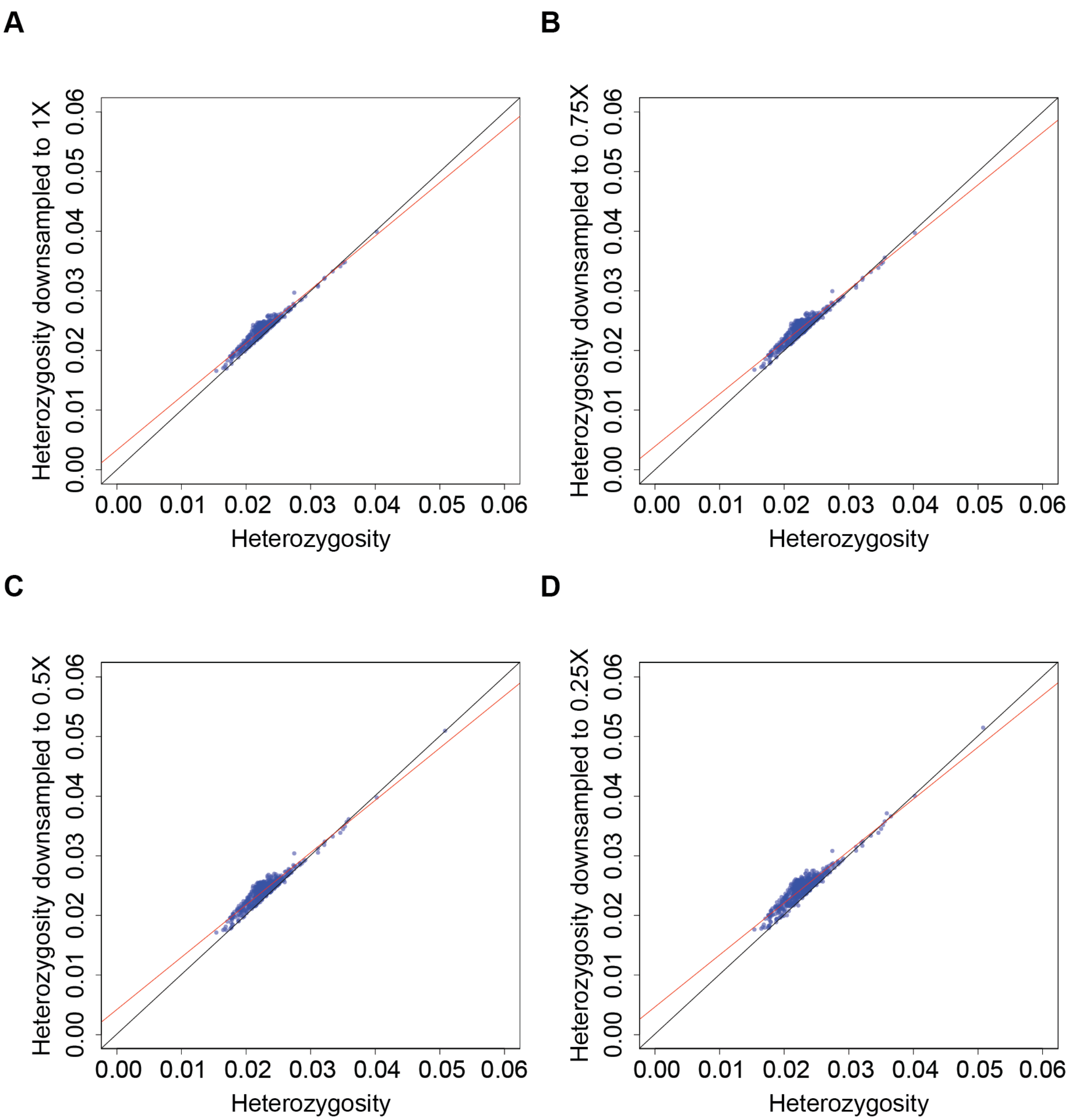


**Fig. S21. Correlation of heterozygosity using all reads per sample vs. downsampled to (A) 1X sequencing depth (Adjusted R-squared: 0.9255, p-value: < 2.2e-16), (B) 0.75X sequencing depth (Adjusted R-squared: 0.9101, p-value: < 2.2e-16), (C) 0.5X sequencing depth (Adjusted R-squared: 0.9046, p-value: < 2.2e-16), (D) 0.25X sequencing depth (Adjusted R-squared: 0.8828, p-value: < 2.2e-16). The black line represents a perfect correlation and the red line represents the linear regression line.**

**Table S1.**

Population-level statistics. The estimate of Tajima's D is based on the mean value across all windows in the sliding window analysis. Heterozygosity was estimated for each sample individually and the mean across samples in a population is reported here. Effective population size was estimated based on the nucleotide diversity, assuming a generation time of one year and a mutation rate of 1 x $10^{-8}$ substitutions/site/generation.

| Population | Sample size | Tajima's D | Heterozygosity | Effective population size ($N_e$) |
|---|---|---|---|---|
| Modern West | 34 | -1.396072 | 0.02332 | 1,138,551 |
| Historical West | 15 | -1.488506 | 0.02478 | 1,048,179 |
| Modern Admixed | 50 | -1.434849 | 0.02302 | 1,197,444 |
| Historical Admixed | 20 | -1.571675 | 0.02903 | 1,155,252 |
| Modern South | 22 | -1.037859 | 0.02293 | 857,481 |
| Historical South | 11 | -1.133698 | 0.02337 | 782,524 |
| Modern East | 71 | -1.260844 | 0.02237 | 1,083,989 |
| Historical East | 46 | -1.679911 | 0.02447 | 1,236,892 |
| Modern Europe | 170 | -1.394707 | 0.02181 | 1,243,678 |
| Historical Europe | 211 | -1.904119 | 0.02273 | 1,788,097 |

**Data S1.**
Significantly enriched GO terms in Fst-outlier windows.

**Data S2.**
Top outlier SNPs (Z>100) located in gene regions.

**Data S3.**
Prevalence of plant pathogens in the different ranges and time periods. Ranges that have a significantly higher prevalence (Welch two sample t-test p-value < 0.05) within a time period are marked in yellow.

**Data S4.**
Overview of herbarium and contemporary *Ambrosia artemisiifolia* samples and sample preparation information used in this study. Abbreviations of source herbaria: B: Botanischer Garten und Botanosches Museum Berlin, Germany; BR: Meise Botanic Garden, Belgium; BRNU: Masaryk University, Czech Republic; C: University of Copenhagen, Denmark; FI: Natural History Museum Firenze, Italy; G: Conservatoire et Jardin botaniques de la Ville de Genève, Switzerland; GH: Harward University, USA; GOET: Universität Göttingen, Germany; GZU: Karl-Franzens-Universität Graz, Austria; HBG: University of Hamburg, Germany; I_ Alexander Ioan Cuza University, Romania; University of Agricultural Sciences and Veterinary Medicine "Ion Ionescu de la Brad", Romania; JE: Friedrich Schiller Universität Jena, Germany; L: Naturalis Leiden, Netherlands; LD: Lund University, Sweden; LY: Université Claude Bernard Lyon, France; MARS: Aix-Marseille Université, France; MO: Missouri Botanical Garden, USA; MPU: Université de Montpellier, France; NEBC: New England Botanical Club, USA; NEU: Université de Neuchâtel, Switzerland; NY: New York Botanical Garden, USA; P: Muséum National d'Histoire Naturelle, France; PH: Academy of Natural Science, USA; PR: National Museum Prague, Czech Republic; PRA: Insitute of Botany, Academy of Sciences, Pruhonice, Czech Republic; PRC: Charles University Prague, Czech Republic; QFA: Université Laval Québec, Canada; ROZ: Sterdoceské Muzeum Roztoky, Czech Republic; S: Swedish Museum of Natural History, Sweden; STU: Staatliches Museum für Naturkunde Stuttgard, Germany; TRH: Norwegian University of Science and Technology, Norway; UPS: Museum of Evolution Lund, Sweden; US: Smithonian Institution, USA; W: Naturhistorisches Museum Wien, Austria; W: Universität Wien, Austria. For samples grown from seeds, location and collection date represent that of the seeds.

**Data S5.**
Overview of outgroup samples from different *Ambrosia* species used to identify putative hybrids and misidentifications in the *Ambrosia artemisiifoila* dataset and their sample preparation methods. For all samples, leaf material from herbarium specimens were used for DNA extraction. Abbreviations of source herbaria: A: Harvard University, USA; GH: Gray Herbarium, Harvard University herbaria, USA; H: Finnish Museum of Natural History, Finland; MASS: University of Massachusetts, USA; NYB: New York Botanical Garden, USA; P: Muséum National d'Histoire Naturelle Paris, France; RBGE: Royal Botanical Garden Edinburgh, UK; S: Swedish Museum of Natural History, Sweden; UC: University Herbarium, University of California, Berkeley, USA.
